# Mechanics of the Spatiotemporal Evolution of Sulcal Pits in the Folding Brain

**DOI:** 10.1101/2025.02.19.639052

**Authors:** Akbar Solhtalab, Yanchen Guo, Ali Gholipour, Weiying Dai, Mir Jalil Razavi

**Author notes:** These authors contributed equally to this work (Akbar Solhtalab, Yanchen Guo).

## Abstract

Understanding the development of complex brain surface morphologies during the fetal stage is essential for uncovering mechanisms behind brain disorders linked to abnormal cortical folding. However, knowledge of the spatiotemporal evolution of fetal brain landmarks is limited due to the lack of longitudinal data capturing multiple timepoints for individual brains. In this study, we develop and validate a true-scale, image-based mechanical model to explore the spatiotemporal evolution of brain sulcal pits in individual fetal brains. Our model, constructed using magnetic resonance imaging (MRI) scans from the first timepoint of longitudinal data, predicts the brain’s surface morphology by comparing the distribution of sulcal pits between predicted models and MRI scans from a later timepoint. This dynamic model elucidates how a smooth fetal brain with primary folds evolves to form secondary and tertiary folds. Our results align with imaging data, showing that sulcal pits are stable during brain development and can serve as key markers linking prenatal and postnatal brain characteristics. The model provides a robust platform to study the evolution of sulcal pits in both healthy and disordered brains, which is crucial as altered sulcal pits patterns are seen in disorders such as autism spectrum disorder (ASD), polymicrogyria, down syndrome, and agenesis of the corpus callosum. This research represents a significant advancement in understanding fetal brain development and its connection to disorders that manifest as abnormal sulcal pit patterns later in life.

## Introduction

Cortical folding, also known as gyrification, is a complex process through which the brain develops convolutions on the surface of the cerebral cortex^1–4^. This folding results in the formation of gyri (ridges) and sulci (valleys), significantly increasing the surface area of the cortex and enhancing cognitive functions as well as efficient information processing^5^. Cortical folding plays a vital role in the cognitive abilities of the human brain^6,7^. Abnormalities in this process have been linked to various neurodevelopmental and psychiatric disorders, which underscores its importance in understanding brain development and function^8–10^. Typically, cortical folding begins during the third trimester of gestation and continues to develop after birth. Initially, primary folds form between 20 and 25 gestational weeks, followed by the emergence of secondary folds between 34 and 36 weeks, and tertiary folds that develop after 36 weeks on the surface of the brain^11^. Primary folds, which are primarily influenced by genetic factors, tend to be more stable than secondary and tertiary folds, which exhibit greater variability across individual brains^12^.

The variability in folding morphologies among individual brains makes it challenging to study the underlying mechanisms of cortical folding. However, recent brain imaging studies^13^ have highlighted that “*sulcal pits*”^14–16^ have the potential to serve as valuable indicators for studying cortical maturation and detecting altered cortical development during the early stages of neurodevelopment. Sulcal pits, which are the first deep sulcal folds, are under strong spatiotemporal genetic control and their spatial distribution exhibits a relatively stable pattern from infancy^17^. Emerging first in utero and reaching their greatest depth in the adult brain, sulcal pits are remarkable anatomical features closely associated with the cytoarchitectonic protomap and human brain function^4,15,18–23^. Evidence indicates that, at term birth, the consistent spatial distributions of sulcal pits in major sulci across individuals are already established and remain relatively stable during the first two years of life^17,24^. Notably, these sulcal pits are genetically controlled during early fetal development^25–28^, their depths have a heritable basis^29^, and sulcal graphs exhibit greater similarity in twin pairs compared to unrelated individuals^30^. Therefore, connectivity, and brain function simultaneously, while mitigating challenges associated with individual variability in folding patterns. In postnatal brains with cerebral malformations, sulcal patterns differ significantly from those in normal brains, both in location and depth^31^. The timing of sulcal pit appearance is positively correlated with heritability estimates, with recent findings suggesting that these estimates for sulcal pits decrease linearly over the evolutionary timeline of gyrification^32^. This suggests that cortical folds formed earlier in the gyrification process are influenced more strongly by genetic factors than those formed later. However, much like the surface morphology of cortical folds, the spatiotemporal evolution of sulcal pits and their stability within individual brains remain poorly understood due to a lack of sufficient longitudinal imaging data of individual brains^13^. Consequently, despite significant insights from imaging studies, the mechanics underlying the formation and evolution of sulcal pits are still not well understood.

Numerous studies have demonstrated the interplay between genetically guided biological processes and physical forces in orchestrating cortical growth and folding^33–37^. As emerging cortical layers develop, variations in cellular density and differential growth rates create mechanical forces that drive the convolution and folding of the cerebral cortex^38^. The differential tangential growth (DTG) theory posits that the faster growth of outer brain layers compared to the slower growth of inner layers initiates mechanical instability, which ultimately leads to cortical folding^39^. This leading hypothesis is supported by several experimental and computational studies^40–49^. In addition to the DTG theory, other factors such as radial constraint^50^, cranial constraint^51^, and axonal tension^52–57^ have also been proposed to play roles in the formation and modulation of cortical folds during brain development.

To date, many biomechanical models developed to study brain folding have primarily relied on simplified and idealized shapes or focused solely on adjusting individual parameters to investigate the growth and folding of the human brain^4,49,58–67^. Moreover, there has been a lack of opportunities to compare the surface morphologies produced by these models with the intricate complexities observed in actual individual brain anatomies. While these models contribute to our basic understanding of brain folding mechanics, they fall short of providing a comprehensive representation of the true-scale nature of brain growth and folding. Therefore, a robust and comprehensive model is needed to elucidate the underlying mechanics of brain folding and to effectively capture the complex evolution of brain surface morphology and sulcal pits.

In this study, we develop true-scale, image-based models that simulate the growth and folding of the brain, shedding light on the emergence and evolution of cortical folds and sulcal pits. We synergistically extend our analysis by integrating real magnetic resonance imaging (MRI) data. Recent advancements in super-resolution reconstruction techniques applied to T2-weighted fetal brain structural MRI^68–72^, fetal brain MRI atlases^73,74^, and automatic segmentation tools^73,75–77^ allow for precise construction and delineation of brain architecture, making it possible to create a true-scale mechanical growth model. We acquired fetal brain MRIs at two time points for each case: one capturing primary folding patterns and the other depicting developed folding patterns. Unlike other studies, the longitudinal images provided a unique opportunity to assess the performance of the mechanical models in predicting the surface morphology of a developing brain. This study advances our understanding of the mechanics of brain gyrification and the spatiotemporal evolution of sulcal pits, which is crucial for deciphering the mechanisms underlying various neurodevelopmental disorders. Additionally, it sheds light on the mechanical mechanisms driving the variability and regularity of cortical folds in the human brain.

## Methods

The data for this study were collected from fetal MRI scans of pregnant women who underwent research fetal MRIs at Boston Children’s Hospital. The study was approved by the institutional review board committee and written informed consent was obtained from all participants. All images were acquired with 3-Tesla Siemens Skyra, Trio, or Prisma scanners using 18- or 30-channel body matrix coils via repeated T2-weighted half-Fourier acquisition single-shot fast spin echo scans in prescribed orthogonal planes of the fetal brain. The slice thickness was 2 mm with no inter-slice gap, in-plane resolution was between 0.9 mm and 1.1 mm, and acquisition matrix size was 256×204, 256×256, or 320×320. Volumetric images were reconstructed using one of the iterative slice-to-volume reconstruction algorithms^69–71^, and the fetal brain was extracted and registered to a standard atlas space in a procedure described in Gholipour et al.^73^. The resulting 3D images had isotropic voxels with a size of 0.8 mm.

In this study, we used super-resolution reconstructed structural (T2-weighted) MRI scans of 24 fetuses, each scanned at two different time points: the first time corresponding to early gestational weeks (GW) (26 to 31 weeks, referred to as the early stage) and the second capturing later gestational weeks (35 to 39 weeks, referred to as the later stage). A detailed overview of the cases and their gestational ages is presented in Table 1.

**Table 1.** Gestational weeks (GW) of two time-point MRI scans from 24 cases.

| Case Number | GW of Scan 1 | GW of Scan 2 | Case Number | GW of Scan 1 | GW of Scan 2 |
| --- | --- | --- | --- | --- | --- |
| 1 | 30.14 | 38.14 | 13 | 26.71 | 38 |
| 2 | 27.14 | 36.86 | 14 | 29.86 | 38.71 |
| 3 | 29.43 | 37 | 15 | 27.86 | 35.86 |
| 4 | 26.71 | 38.71 | 16 | 29.71 | 36.29 |
| 5 | 30 | 37.86 | 17 | 27.71 | 36 |
| 6 | 28.14 | 37.14 | 18 | 28.75 | 37 |
| 7 | 28.86 | 36.29 | 19 | 29.57 | 39.57 |
| 8 | 29.57 | 35.57 | 20 | 29.43 | 39.57 |
| 9 | 31.86 | 35.57 | 21 | 27.71 | 36.43 |
| 10 | 26.78 | 35.47 | 22 | 28.71 | 36.71 |
| 11 | 28 | 33 | 23 | 29 | 35.57 |
| 12 | 28 | 38 | 24 | 27.71 | 34 |

### Reconstruction of white matter and pial surfaces of early-stage scans

Extracting cortical and white matter surfaces from MRI data is a critical step in constructing accurate finite element method (FEM) models of the entire brain. To develop a true-scale mechanical model for brain growth and folding, we reconstructed the initial smooth state of fetal brain scans obtained during the early stages of development (first time point scans) (Fig. 1a). We used a combination of atlas-based segmentation and deep learning^73,75^ to segment the super-resolution reconstructed fetal MRI scans to different tissue types including the developing white matter and the cortical plate (the cortical gray matter). However, the pial surface from automatic gray matter segmentation maps often contained some structural errors, such as holes and undesired attached regions (Fig. 1b). To improve the accuracy of the extraction, we manually refined the segmented gray matter surface and white matter surface using the 3D Slicer software (Fig. 1c). This manual adjustment was important for correcting the intricacies and details that automatic thresholding methods might miss, thereby ensuring a precise representation of cortical surface for further analysis.

**Fig. 1.**
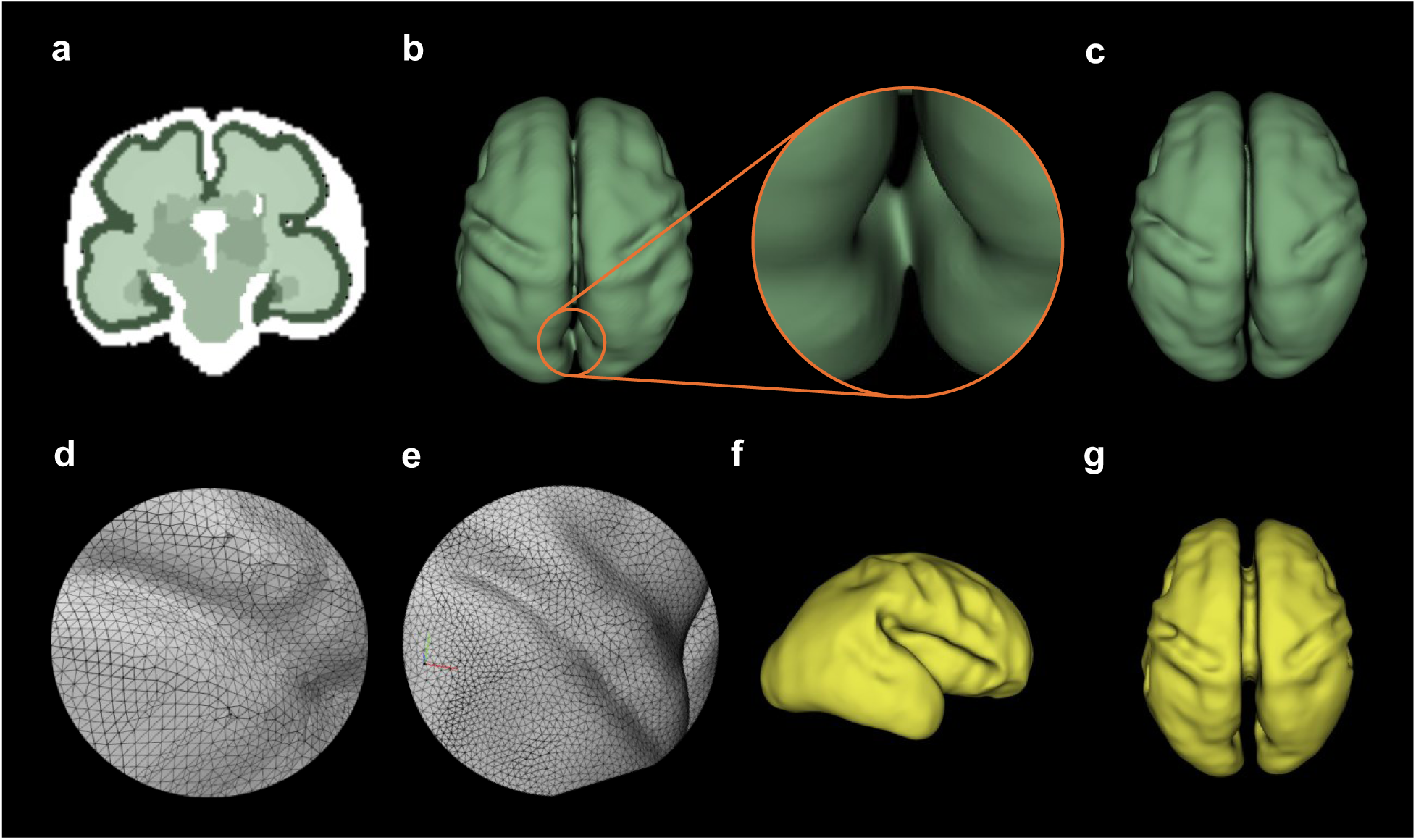
Reconstruction of pial and white matter surfaces. **a** MRI image of a case at 27 gestational weeks. **b** Pial surface from automatic gray matter segmentation without manual modification, showing a close-up of a structural error. **c** Pial surface after manual modification. (d) Irregular mesh of the surface. **e** Surface after smoothing and uniform remeshing, **f-g** Sagittal and axial views of the white matter surface.

The cortical surface extraction process produced an irregular triangular mesh containing approximately 300,000 vertices (Fig. 1d). To enhance the mesh’s uniformity and optimize it for further analyses, we applied Laplacian and Tubin smoothing techniques. These methods improved the signal-to-noise ratio by reducing irregularities that arose during the surface reconstruction from MR images^78^. Following smoothing, we performed uniform remeshing of the surface (Fig. 1e), resulting in refined meshes that serve as the foundation for subsequent quantitative surface measurements. The precision of these results depends heavily on the mesh quality. Through this refinement, we aimed to create a more reliable and consistent cortical surface representation, ensuring an optimal mesh for robust FEM simulations of the brain.

The selection of smoothing iteration values was vital for achieving the desired results. After careful experimentation, we found that 50 iterations of Laplacian smoothing and 100 iterations of Tubin smoothing yielded optimal outcomes, effectively preventing over-smoothing and excessive shrinkage of the cortical surface^79^. These settings significantly improved the signal-to-noise ratio by reducing reconstruction noise, resulting in a more uniform, simplified, and smoothed surface. To generate the white matter surface from the gray matter surface, we applied an inward offset of 1.4 *mm*, corresponding to the cortical thickness (Fig. 1f-1g). This approach was chosen because extracting the white matter surface separately often leads to challenges in aligning the nodes at the gray–white matter interface. Separate extraction typically results in mismatched surfaces and discontinuities at the interface. This assumption aligns with findings from imaging studies and previous computational models, which show that cortical thickness is largely uniform across the brain prior to folding, when the brain surface is still smooth^33,80^.

### Reconstruction of white matter surfaces of later-stage scans

To perform the geometric measurements and extract the sulcal and gyral peaks, we focused on the white matter surfaces corresponding to the second time point, representing the later gestational weeks as outlined in Table 1. The extraction process followed the procedures described in the previous section. Initially, the white matter segmentation map (Fig. 2a) was generated automatically; however, the white matter surface derived from this map exhibited several geometric inaccuracies (Fig. 2b). To address these issues, manual modifications were made to correct the structural errors. Subsequently, smoothing and remeshing procedures were applied to refine the surface. The resulting map, after manual correction, smoothing, and remeshing, is shown in both axial and sagittal views (Fig. 2c-2d). These steps were essential for ensuring that the white matter surfaces were of sufficient quality for accurate geometric measurements and further analysis.

**Fig. 2.**
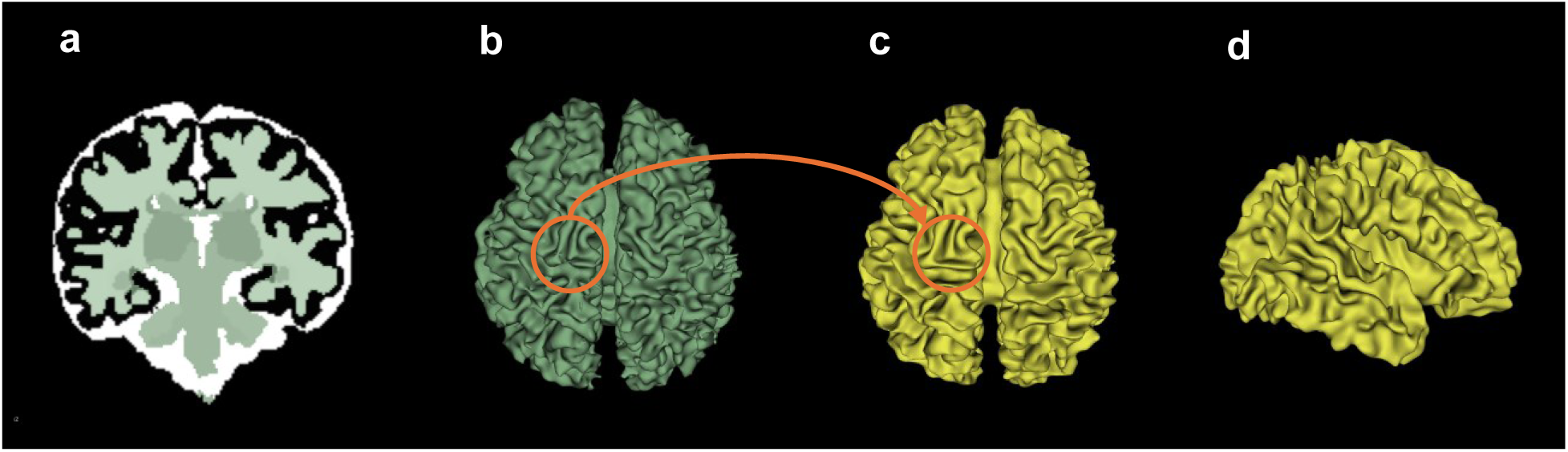
Reconstruction of the white matter surface from the second time point (later-stage scan). **a** Automatic segmentation of the original MRI image at 38 gestational weeks, **b** The extracted white matter surface without manual modification, **c-d** Axial and sagittal views of the white matter surface following manual modification, smoothing, and remeshing.

### Constitutive Framework

To model the growth and folding of the human brain, we adopt the well-established DTG mechanism as the primary driver of brain folding^39^. In DTG, differential expansion rates between the rapidly growing cortical plate and the slower-expanding subcortical regions generate compressive forces. These forces create mechanical instability, ultimately leading to the buckling of the cortical plate, which drives the characteristic folding patterns of the brain^39^. Since the cortex and white matter exhibit distinct growth behaviors, it is crucial to implement unique constitutive equations for each region to accurately capture their individual growth mechanisms.

#### Multiplicative decomposition of the deformation gradient

We consider a continuum body, denoted as *B*_*R*_, which is defined by the region of space it occupies in a fixed reference configuration. Within this body, arbitrary material points are represented as **X**. As time progresses, the referential body *B*_*R*_ experiences a motion described by **x** = *φ*(**X**, *t*), which leads to the formation of the current deformed body, denoted as *B*_*t*_. The deformation gradient deformation gradient

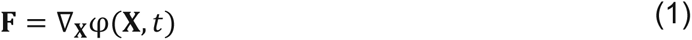

captures the transformation undergone by the body during this process. The symbol ∇ represents the gradient operator applied with respect to the material point **X** in the reference configuration.

In the field of continuum mechanics, the noticeable deformation of biological tissue can be interpreted as a combination of two mechanisms: growth, the deformation due to increase in size or number of cells and cell processes, and elastic deformation, which is caused by mechanical forces^81^. Then, the total deformation gradient can be expressed as follows

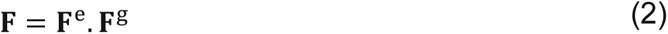

where **F**^e^is the elastic deformation gradient tensor and **F**^g^ is the growth deformation gradient tensor. This decomposition enables us to account for both elastic **F**^e^ and irreversible growth-related **F**^g^components, which provide a comprehensive understanding of the overall deformation experienced by the body.

#### Constitutive model for cortex

The imaging data indicate that cortical thickness undergoes changes during the early developmental stage^82^. However, in the later stages, it is primarily the alteration in surface area that initiates cortical folding. These findings suggest that the growth of the cortex primarily occurs tangentially, as supported by several studies^58,83^. Therefore, we adopted a model for cortical growth that considers in-plane area expansion, while it assumes purely elastic behavior in the direction normal to the cortical layer, following the approach proposed by Holland et al.^57^

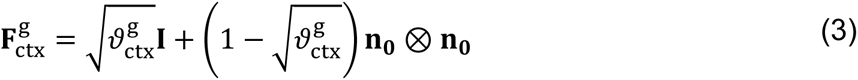

Eq. (3) incorporates a growth multiplier 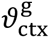, the referential unit normal **n_0_** of the pial surface, second order unit tensor **I**, and accounts for the increase in the cortical area. The growth multiplier 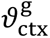 can be expressed as follows

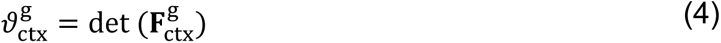

A linear kinetic model for the growth of the cortex (gray matter) was used^57^

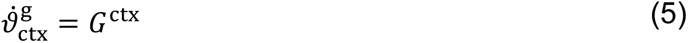

where *G*^ctx^ is the cortical growth rate. Then the elastic deformation gradient can be explicitly calculated by the following equation

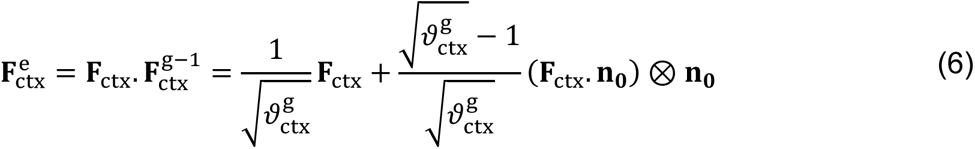

Lastly, the elastic left (**b**^e^) and right (**C**^e^) Cauchy-Green tensors can be derived from 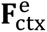, respectively

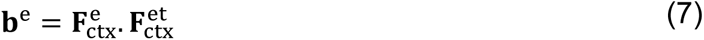

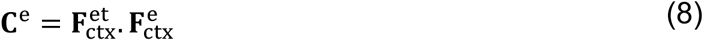

#### Constitutive model for white matter

During the developmental process, there is an observed increase in the volume of white matter^57^. Therefore, we consider that the growth of the white matter is isotropic. The growth tensor is

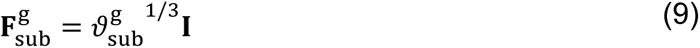

where *I* denotes the second order unit tensor and 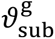 is the growth parameter represents the increase in the volume of the white matter

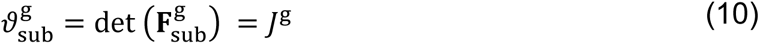

The growth rate of the white matter is defined as follows:

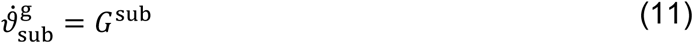

Similar to the previous brain folding models^40,42^, we used a standard neo-Hookean hyperelastic material model with the depicted free energy function in Eq. 12 for the white matter and the cortex

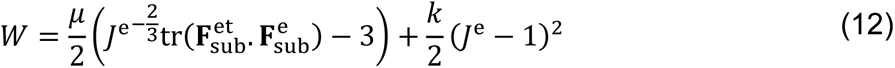

where *μ* is the shear modulus, *k* is the bulk modulus, and 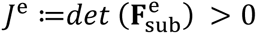 is the Jacobian. We introduce the Cauchy stress tensor of the white matter generated by the elastic deformation tensor in the material configuration as follows

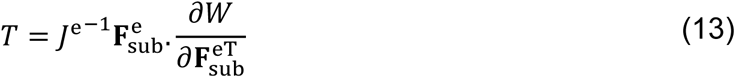

where *j*^e^ is the elastic volume change ratio.

### Computational Model

The simulation geometry is based on T2-weighted, motion-corrected MRI scans of fetal brains at the early-stage of development, as detailed in Table 1. Using the methods described in previous sections, we constructed the cortical layer and white matter from the MRI data. The two volumes were merged and uniformly meshed with a mesh size of 0.4 mm, resulting in approximately 7 million tetrahedrons. This meshing ensured a minimum of four tetrahedral layers across the cortical thickness. The constitutive models described earlier were implemented in the Abaqus/Explicit finite element package via a user-defined material subroutine (VUMAT). We modeled the brain as a neo-Hookean material with the strain energy density defined by Eq. (12). The majority of experimental evidence supports a ratio of 1 to 2 between cortex and white matter stiffness in the human brain^84–86^. We considered the same shear modulus for both the cortex and white matter^59^. A uniform pressure of 0.7*μ* (*μ* is the shear modulus of gray matter) is applied to the cortical surface to simulate the effect of the meninges and skull, resulting in a flattened sulcal morphology^40^. To simulate brain folding, we specified a tangential growth rate for the cortical layer and an isotropic growth rate for the white matter with growth ratio of *G*^ctx^/*G*^sub^= 4. The differential growth rates between the cortical layer and white matter drive the expansion and gyrification process.

### Sulcal Pits Extraction and Analysis

Here, we delineate the process for analyzing sulcal pits, which includes brain registration, sulcal pits extraction, and the matching of sulcal pits between two brains. The similarity of sulcal pits from the two brains is computed and subsequently analyzed.

#### Sulcal pits extraction

Sulcal pits are defined as the deepest points of the sulcal basins of the cerebral cortex^15,16^. We extracted the sulcal pits automatically according to the pipeline described in Auzias et al. ^24^. The extraction process primarily involves two steps: estimating a sulcal depth map and extracting the pits using a watershed-by-flooding algorithm^87^, while filtering out noisy sulcal pits. The Depth Potential Function (DPF) introduced in Boucher et al.^88^ was used to estimate the sulcal depth map. The DPF is reference free, independent of brain size and it has only one parameter *α* = 0.03 that controls the trade-off between mean curvature and average convexity. After estimating the sulcal depth map using DPF, we applied the watershed-by-flooding algorithm to generate sulcal basins and the corresponding sulcal pits. During the watershed flooding, sulcal basins were merged if ridge height (R) was less than Threshold_R = 1.5 and the distance between two pits (D) was less than Threshold_D = 20. At the end of the flooding, any sulcal basin with a basin area (A) less than Threshold_A = 50 was further merged with its neighbor that shares the longest border. Figure 3 illustrates the extracted sulcal pits from early-stage fetal brain MRI, alongside FEM-simulated brains at seven developmental stages, presented in chronological order. For comparison, the sulcal pits from the later-stage brain MRI from the same fetus are also included in Fig. 3.

**Fig. 3.**
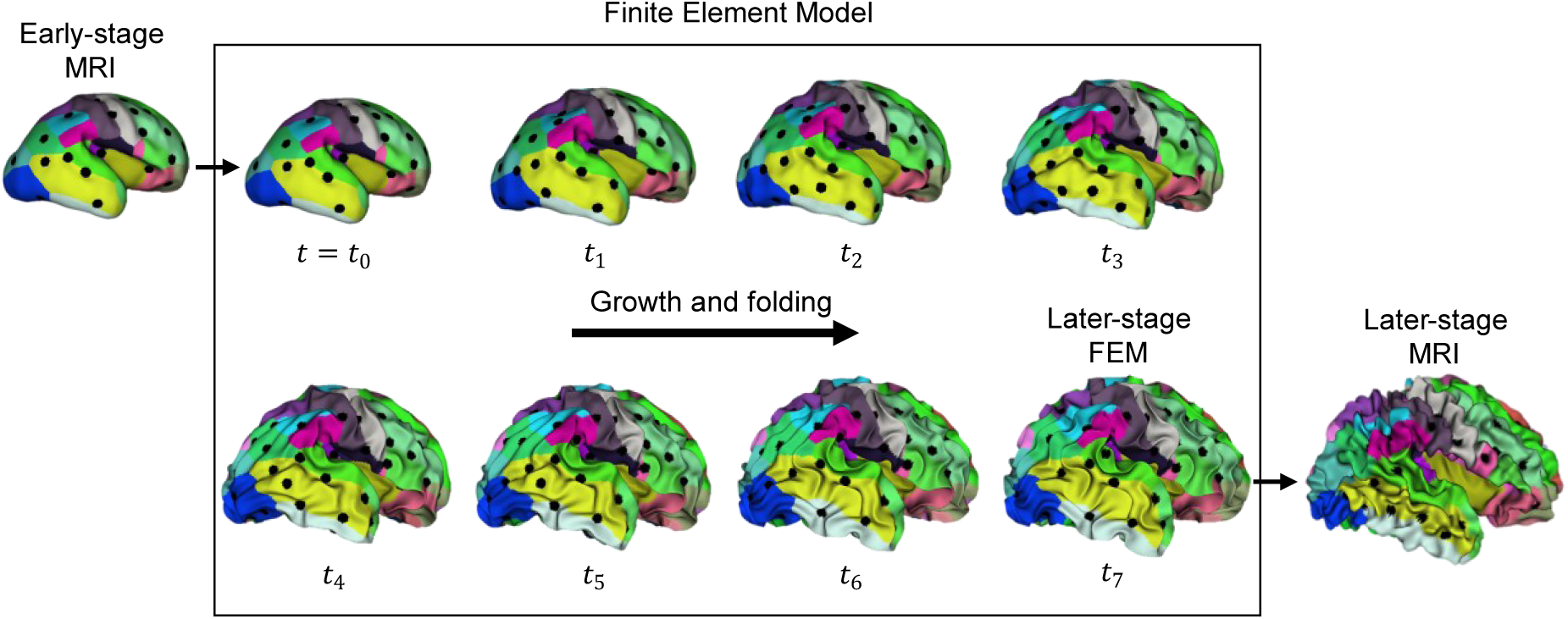
Extracted sulcal pits overlaid on the corresponding brain white matter surface partition map for various developmental stages. The left panel displays sulcal pits from the early-stage MRI, while the middle panel shows the brain model at the initial stage (t_0)_, serving as the input for the FEM model, along with the FEM-simulated brain models at seven developmental stages (t_1 t_o t_7)_. The right panel presents the later-stage MRI. Note that the brain model at the initial stage is identical to that from the early-stage MRI.

#### Sulcal pits matching

The pattern of sulcal pits extracted from each brain was represented as a 3D point cloud. Consequently, the task of matching sulcal pits between two brains was thus formulated as a points-matching problem involving these two 3D point clouds. To solve this, we employed the Reweighted Random Walk Matching (RRWM) algorithm^89^, which was used to perform point cloud matching between two brains by aligning their respective sulcal pits. For each brain, we considered the 3D coordinates [*x*, *y*, *z*] of the sulcal pits as the geometric features of the points. The goal was to find the optimal correspondence between the sulcal pits of two brains by minimizing the differences in these features. Let *P* ∈ *R*^*I*×3^ and *Q* ∈ *R*^*j*×3^ represent the two sulcal pit point clouds, with *I* and *j* being the number of sulcal pits in each brain. We first constructed an affinity matrix *A* ∈ *R*^*I*×*j*^ using a Gaussian kernel, which quantified the similarity between potential correspondences of points in *P* and *Q* as presented in Eq. 14

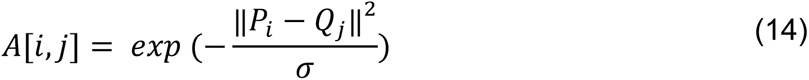

where *P*_*i*_ and *Q*_*j*_ are the *i*th row of P and *j*th row of Q, *i* = 1, ⋯, *I*, *j* = 1, ⋯ *j*. ‖*P*_*i*_ − *Q*_*j*_‖^2^ is the squared Euclidean distance between *P*_*i*_ and *Q*_*j*_. σ is a scaling parameter that controls the “spread” of the aussian function, and it is set to 100.

The RRWM algorithm was then applied to the affinity matrix *A*, yielding a matching matrix *M* ∈ *R*^*I*×*j*^, where each entry represented the likelihood of correspondence between a pair of sulcal pits from the two 3D point clouds. After discretizing the matching matrix *M*, we obtained a set of matched sulcal pits, with the number of matches being the minimum of *I* and *j*. For each pair of matched sulcal pits, we further calculated a pairwise similarity score using the Gaussian kernel affinity function, resulting in similarity values ranging from 0 to 1. To refine the matches, we applied a similarity threshold of 0.5, eliminating any weakly paired sulcal pit pairs. Only strongly paired sulcal pits are considered as paired sulcal pits.

#### Brain registration

Since our primary focus is on matching sulcal pits between two brains of different sizes, such as early-stage and later-stage brains, it is important to consider that brain size, which influences the 3D positions [*x*, *y*, *z*] of the sulcal pits, can affect the accuracy of the point cloud matching algorithm. To enhance the robustness of sulcal pits matching results, we designed a two-stage position-based registration: the first stage coarse registration, sulcal pits matching, second stage refined registration, sulcal pits matching again.

The first stage involved a coarse registration by rescaling the early-stage brain along the positive and negative x, y, z dimensions to align it with the reference brain from the later stage. The coarse registration step works well because the origins of both brains are located at the same anatomical position. This rescaling was necessary because the growth rates of the brain differ across these six dimensions. The rescaling factors were determined based on the relative positions of the sulcal pits in corresponding space of the two brains. For instance, let *P*_*xp*_ ∈ *R*^*N*×3^ represent N sulcal pits located in the positive x-dimension of the brain to be scaled, and *Q*_*xp*_ ∈ *R*^*M*×3^ represent M sulcal pits in the positive x-dimension of the reference brain. The positive x-factor *f*_*xp*_ was calculated by Eq. 15 as follows.

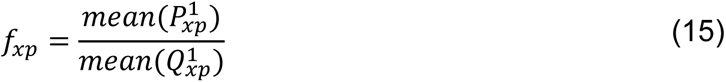

where 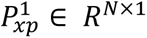 denotes the x-coordinates of the N sulcal pits from the brain being scaled and 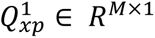 denotes the x-coordinates of the M sulcal pits from the reference brain. From the two brains with a coarse registration, we can identify strongly paired sulcal pits (with similarity scores exceeding 0.5). as described in the Sulcal pits matching section.

The second stage involved a refined linear registration^90^ aiming to determine the affine transformation, including translation and rotation, that optimally aligns the strongly paired sulcal pits identified in the first stage. Sulcal pits matching was then performed based on the linearly registered brains, leading to the final matching outcomes. This two-stage sulcal pits matching approach significantly enhances the accuracy and reliability of our results.

#### Similarity degree of sulcal pits (SDSP)

*SDSP* between the two brains was defined based on the matching results from the two-stage method. The average similarity across strongly paired sulcal pits was utilized to quantify the overall SDSP between the two brains.

### Data Analysis

Heatmaps were utilized to illustrate the SDSP between early-stage MRI and later-stage MRI, between later-stage FEM models and later-stage MRI, between early-stage and later-stage FEM models. In each heatmap, the diagonal values represent the similarity within the same subject, while the off-diagonal values reflect similarity between different subjects. Based on the stability of sulcal pits within individual brains, we hypothesized that the diagonal SDSP value would be the largest on the same row. In other words, the distribution of sulcal pits at the early-stage should be more similar to those at the later-stage of the same subject, rather than to those of other subjects. To examine the evolution of sulcal pits during brain development, we used FEM to simulate seven developmental stages (t_1_, t_2_, …, t_7_) starting from the early-stage MRI model (t_0_). We then calculated the SDSP values between these developmental stages and t_0_. Paired t-tests were conducted to compare the SDSP values between two adjacent development stages and the early stage, specifically comparing the SDSP values between t_i_ and t_0_ versus those between t_i-1_ and t_0_ (i =1, …, 7). These t-tests aimed to assess whether there were statistically significant differences in the similarity of sulcal pits between adjacent stages relative to the early stage.

## Results

In this study, we developed a true-scale, image-based mechanical model to investigate the mechanics of brain development and folding, with a focus on the spatiotemporal evolution of sulcal pits (deep sulcal roots). Using our growth framework, we applied the model to 24 fetal brain scans at the early-stage time point and simulated the folding process up to the later-stage time point. Before developing the mechanical model, the longitudinal MRI data were analyzed to assess how many later-stage scans could be accurately associated with their corresponding early-stage scans. Specifically, the goal was to determine the number of individual cases with two MRI scans that exhibit a greater SDSP (similarity degree of sulcal pits) between the early-stage and later-stage time points from the same individuals compared to SDSP from different individuals. This step was crucial for selecting cases as ground truth for developing mechanical models, ensuring that the similarity metric demonstrated a stronger correlation between the two stages. In this section, quantified results from the simulations and the MRI data analysis are presented.

### Analysis of longitudinal MRI data

A total of 24 fetal brains, each with two scans, one at an early stage and another at a later stage of development, were analyzed (Table 1). Figure 4 illustrates the SDSP for all 24 cases, evaluating the similarities between their early- and later-stage MRI scans. We expected the SDSP between two stages of the same fetus to be higher than that between different fetuses. The diagonal SDSP values in Fig. 4 represent the similarity between the early-stage scan and later-stage scan from the same fetus. In contrast, the off-diagonal SDSP values in each row indicate the similarity between the early-stage scan of each case and the later-stage scans of other cases. When the diagonal value is the highest in each row, it suggests that the later-stage scan can be successfully matched with the early-stage scan of the same fetus using our developed sulcal pits similarity algorithm. However, in instances where the diagonal values are not the largest, the algorithm matches the early-stage scan of one case with the later-stage scan of an unrelated case.

**Fig. 4.**
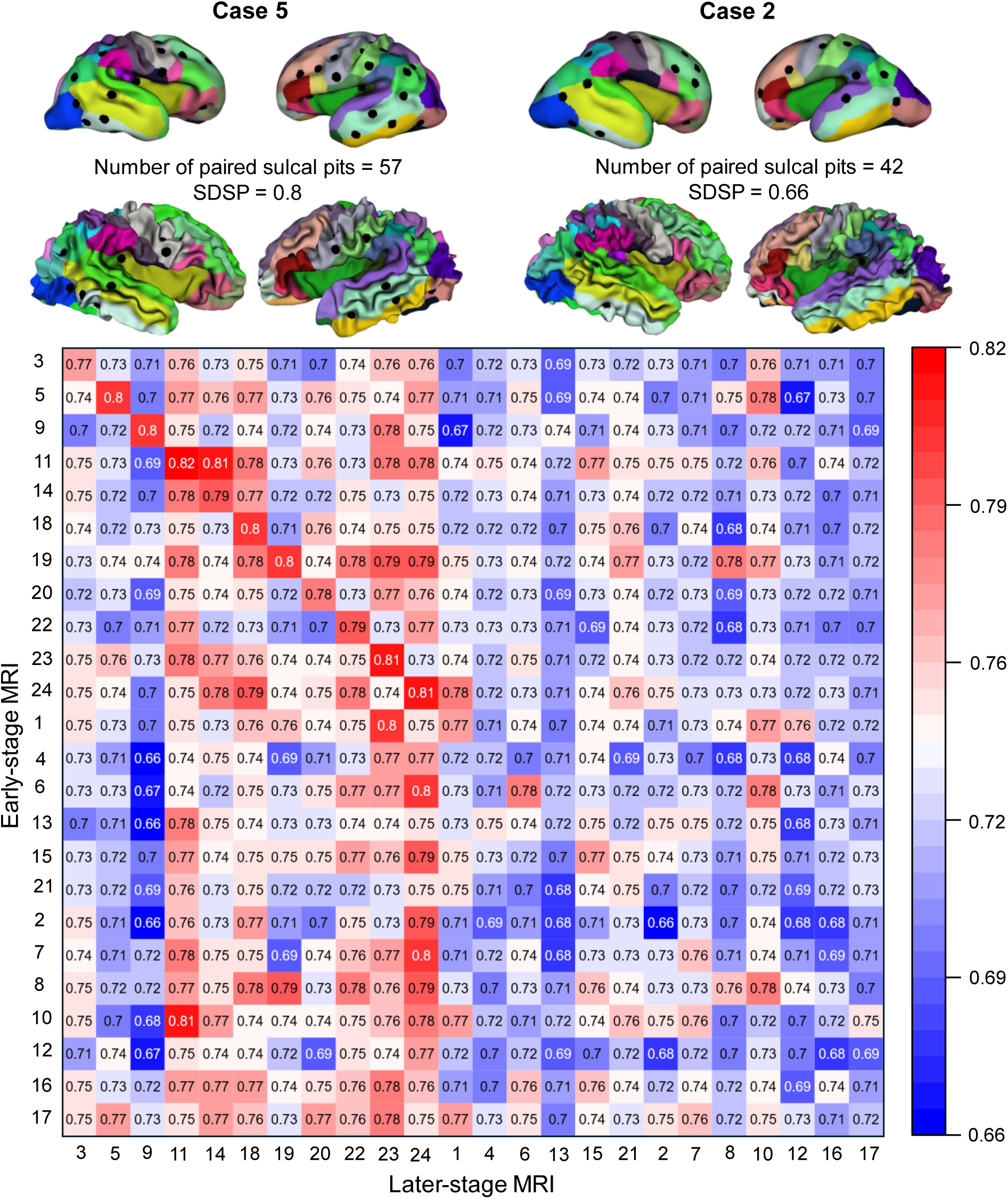
The heatmap illustrates similarity degree of sulcal pits (SDSP) between early-stage and later-stage MRI scans across all subject pairs. The diagonal represents the similarity within the same subject, while the off-diagonal elements show similarity between different subjects. Due to variations in MRI scan quality, only the first 11 subjects exhibit higher similarity with their own developed brain, making them the reliable cases in our ground truth dataset.

The algorithm successfully identified 11 cases (sorted and shown as the first 11 rows in Fig. 4) that exhibited greater similarity between their own early and later-stage scans compared to the scans of other cases. This suggests stability in the formation of primary sulcal pits up to the second time point. In contrast, the remaining 13 cases (shown as the last 13 rows in Fig. 4) showed lower similarity between their own longitudinal scans than between their early-stage scans with the later-stage scans of other cases. To understand the source of this inconsistency, we investigated potential factors such as the effect of time gaps between two longitudinal scans.

Noting that the early-stage and later-stage scans were not all acquired at the same gestational weeks (see Table 1), we examined the diagonal and off-diagonal SDSP values of longitudinal scans in relation to the early-stage gestational weeks, later-stage gestational weeks, and the time gap between longitudinal scans (Fig. 5). The SDSP between early-stage and later-stage MRI scans was positively associated with early-stage (first time point) gestational weeks for diagonal cases (Fig. 5a, p = 0.0052) but was not significant for off-diagonal and diagonal values (Fig. 5a, p = 0.14). It was negatively associated with later-stage (second time point) gestational weeks for both values (Fig. 5b, p = 0.047 and p < 10^-6^, respectively) and negatively associated with time gaps for both values as well (Fig. 5c, p = 6.49 ×10^-4^ and p < 10^-6^, respectively). These relationships are expected, as cortical folding progresses and sulcal pits deepen over time. As a result, the SDSP is smaller with earlier stages of brain development and larger with later stages. The variation in MRI scan times may contribute to the mismatch between the early-stage and later-stage scan from different cases for the last 13 fetuses in Fig. 4. For instance, compared to the early-stage scan of case 4, its own later-stage scan has smaller SDSP than the later-stage scans of cases 23 and 24. This is caused by the smaller GW of case 4 at the early-stage scan (smaller SDSP, red line in Fig. 5a) and smaller GWs of cases 23 and 24 at the later-stage scans (larger SDSP, blue line in Fig. 5b). The fetuses with smaller GWs typically exhibit smoother brain surfaces, characterized by shallower sulcal pits. For smooth brain surfaces of different cases, subtle variations of sulcal pits are expected and therefore larger SDSP values were observed between the early-stage scan of case 4 and later-stage scan of cases 23 and 24. To put together, the timing of these longitudinal MRI scans is the confounding factors in matching them from the same fetuses. Another contributing factor is the limited MRI scan quality of some of the unmatched cases, which impacted the ability to accurately match the later-stage brain MRI scans to their early-stage counterparts based on the similarity of their sulcal pits.

**Fig. 5.**
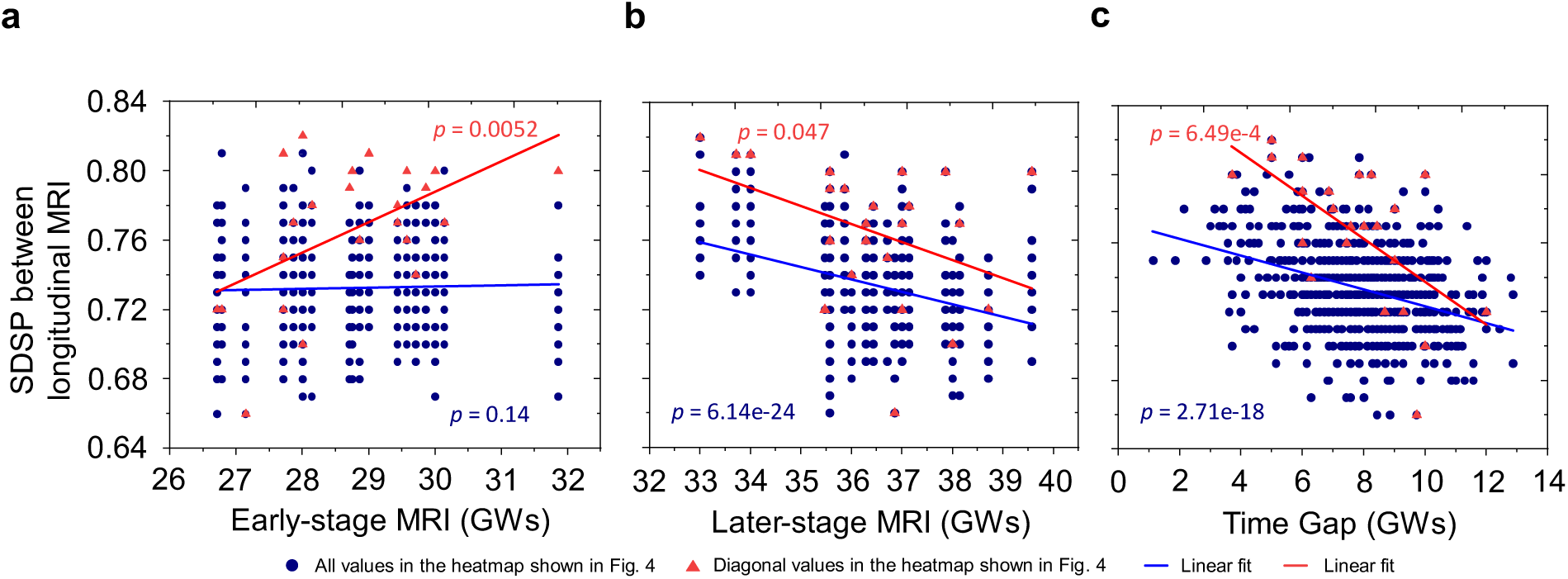
The dependency of the similarity degree of sulcal pits (SDSP) between early-stage and later-stage longitudinal MRI scans on the timing of MRI acquisitions and the time gap between scans. **a** The SDSP between early-stage and later-stage MRI scans is positively correlated with the gestational weeks at the time of the early-stage MRI for diagonal cases (p = 0.0052) and not significantly correlated for off-diagonal values (p = 0.14). **b** The SDSP is negatively associated with later-stage gestational weeks for both diagonal (p = 0.047) and off-diagonal values (p < 10^-6^). **c** The SDSP is negatively associated with time gaps between scans for both diagonal (p=6.49×10^-4^) and off-diagonal values (p<10^-6^). Larger time gaps between longitudinal scans are associated with a decrease in SDSP between early-stage and later-stage MRI scans.

### Growth and folding of the brain model

Figure 6 illustrates the growth and gyrification of the brain model for an individual with MRI scans taken at two time points. The first time point, at 26 GW, was used to construct the initial mechanical model, which at this stage is smooth and contains only a portion of the primary folds (t_0_=0). The model then grows according to the growth framework outlined in the Methods section, gradually developing folds over time, up to the second time point at 38 GW. In alignment with the differential tangential growth (DTG) mechanism, the mismatch in growth rates between the faster-growing cortex (gray matter) and the slower-expanding white matter (subcortex) generates compressive forces in the cortex, ultimately leading to mechanical instability and brain folding. The model successfully captures the emergence of secondary and tertiary folds from the primary folds, as well as illustrates the spatiotemporal evolution of sulcal pits. The folding morphology and sulcal pit patterns observed in the model at the second time point qualitatively align with the folding patterns and sulcal pits seen in the corresponding MRI scans. Figure 6 presents seven time steps of the model’s growth and folding, although the mechanical model is capable of generating numerous time steps for any growth scenario. This ability to simulate multiple stages of growth is a key advantage, as it offers more detailed insights compared to the typically limited number of longitudinal MRI data points (e.g., two or three) available for each individual case. The dynamic growth and folding process of the model is captured in Supplementary Video 1.

**Fig. 6.**
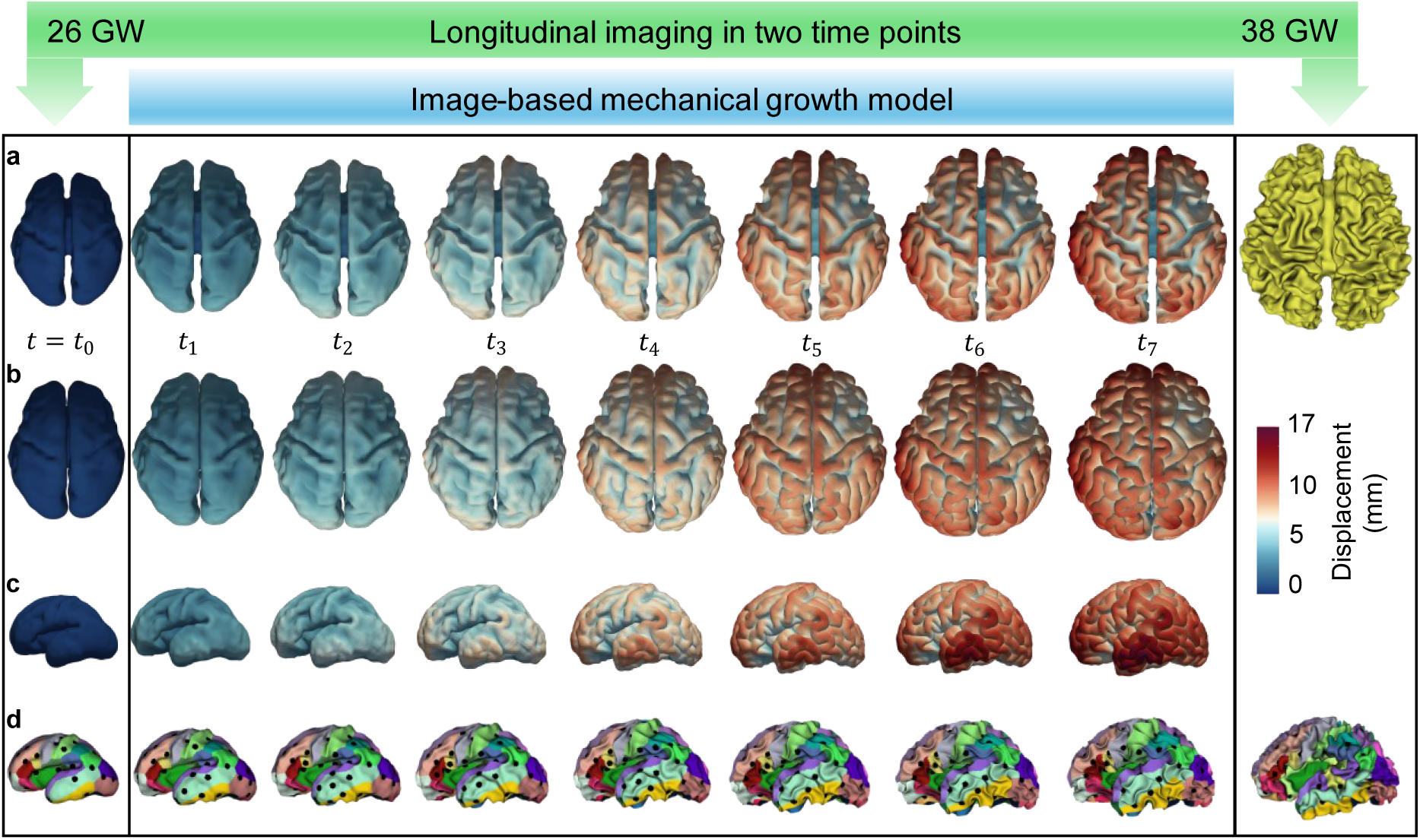
Growth and folding of the simulated brain compared to corresponding MRI-derived white matter and gray matter surfaces at 26 GW and 38 GW. The simulation starts with a smooth fetal brain at approximately 26 GW and demonstrates the development of gyrification driven by the tangential growth of the cortical layer by 38 GW. **a** Left column: white matter surface from fetal MRI at 26 GW. Middle column: white matter surface of the brain model during growth and folding (t_1 t_o t_7)_. Right column: white matter surface from fetal MRI at 38 GW. **b** Left column: gray matter surface from fetal MRI at 26 GW. Middle column: gray matter surface of the brain model during growth and folding (t_1 t_o t_7)_. Right column: gray matter surface from fetal MRI at 38 GW. **c** Left column: side view of the gray matter surface from fetal MRI at 26 GW. Middle column: side view of the gray matter surface of the brain model during growth and folding (t_1 t_o t_7)_. **d** Left column: sulcal pits from the fetal brain at 26 GW extracted and overlaid on the white matter surface from MRI. Middle column: sulcal pits of the brain model during growth and folding (t_1 t_o t_7)_. Right column: sulcal pits extracted and overlaid on the white matter surface from MRI at 38 GW. The color contour represents displacement (in mm) in the model.

### Performance of mechanical models and prediction of sulcal pits evolution

We simulated the growth and folding models for all 24 cases, including the 13 cases where early-stage and later-stage SDSP did not yield the highest values for the same fetuses. Figure 7 presents a heatmap illustrating the similarity of sulcal pits between the later-stage FEM models and the associated later-stage MRI scans for these 24 fetal cases. As expected, the 11 cases that exhibited greatest brain similarity between the early-stage and later-stage MRI scans from the same fetuses (indicated by the largest value in the diagonal element within each row in Fig. 4, rows 1 to 11) also showed highest brain similarity between the later-stage FEM models and MRI scans for the same fetuses (with the largest diagonal similarity in Fig. 7, rows 1 to 11). This result demonstrates the strong predictive performance of the mechanical models in capturing the later-stage brain morphology and sulcal pit patterns when supplied with robust ground-truth data. Interestingly, six cases did not demonstrate the greatest brain similarity between the early-stage and later-stage MRI scans from the same fetuses (see Fig. 4, rows 12 to 17) but did show the highest similarity between the later-stage FEM model and MRI scans from the same fetuses (i.e., largest diagonal similarity in Fig. 7 within the same rows, rows 12 to 17). This finding suggests that the later-stage brain model generated from FEM model shares a more comparable distribution of sulcal pits with the later-stage brain MRI scan for the same fetus. However, seven cases exhibited the largest brain similarity neither between the early-stage and later-stage MRI scans from the same fetuses (as shown in Fig. 4, rows 18 to 24) nor between the later-stage FEM model and MRI scans from the same fetuses (illustrated in Fig. 7, rows 18 to 24). The limited quality of these early-stage brain MRI scans may have hindered the FEM models from achieving a similar distribution of sulcal pits to the later-stage MRI scans. Nonetheless, even in these seven cases, the numbers of off-diagonal elements exceeding the diagonal values was significantly smaller (p = 0.028) in the heatmap between the later-stage FEM and MRI scans (5.57 ± 6.27) than the numbers between the early-stage and later-stage MRI scans (12.29 ± 7.76). In each of these three scenarios, FEM models successfully simulate the growth and folding of a smooth brain, resulting in a folded brain that preserves the sulcal pits distribution observed in the real-MRI scans.

**Fig. 7.**
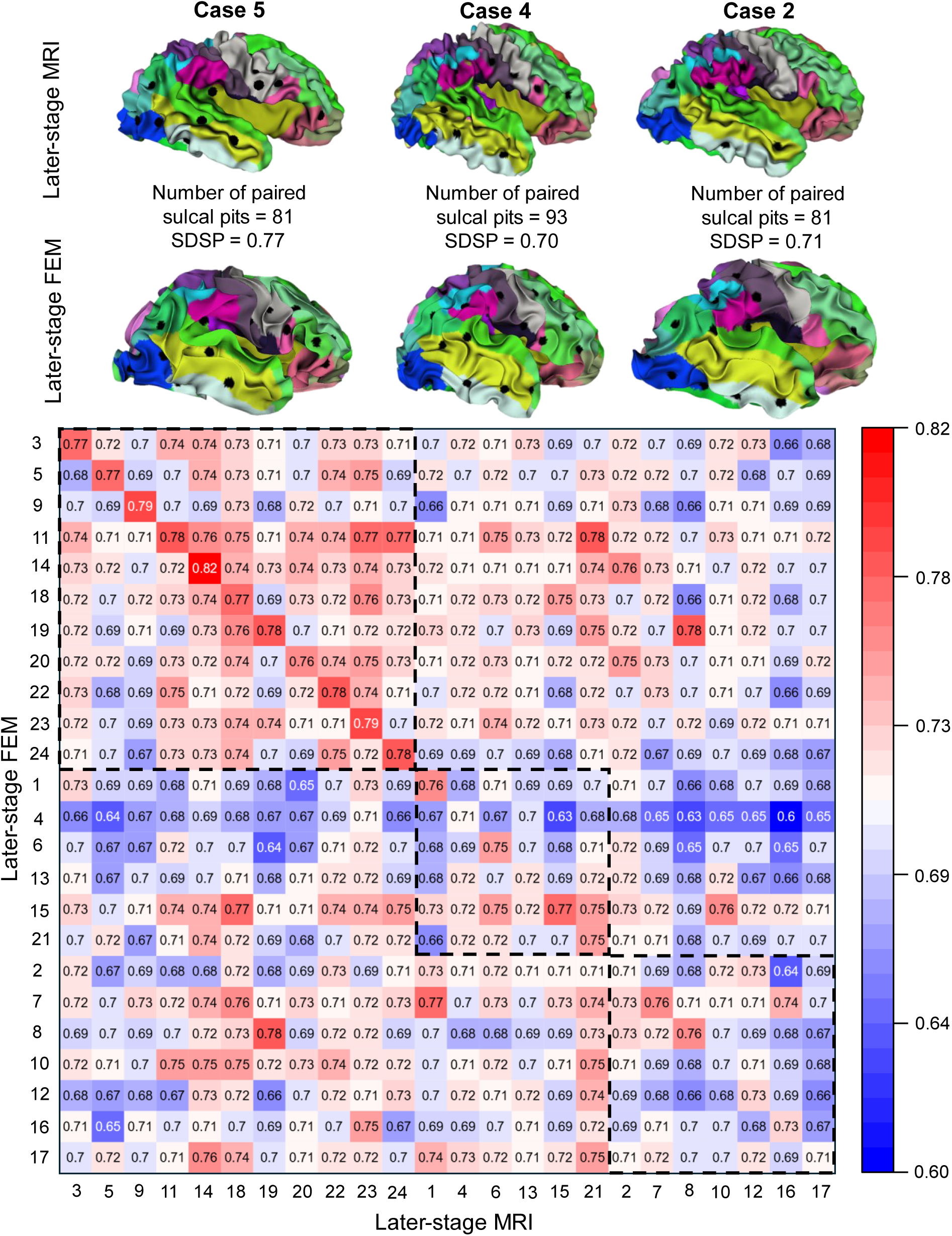
Sulcal pits-based similarity (SDSP) between later-stage FEM models and later-stage MRI scans across all subject pairs. Diagonal values indicate within-subject similarity, while off-diagonal values represent between-subject similarity. Rows 1 to 11 (top-left dashed box): These show higher SDSP values between the later-stage FEM model and their corresponding later-stage MRI scans compared to unrelated cases. Case 5, along with its extracted sulcal pits from the FEM model and MRI scans, is provided as an example of this category. Rows 12 to 17 (middle dashed box): These display higher similarity between the later-stage FEM model and MRI scans from the same fetuses for cases where the greatest similarity was not observed between the early-stage and later-stage MRI scans from the same fetuses. Case 4, with its extracted sulcal pits from the FEM model and MRI scans, is presented as an example of this category. Rows 18 to 24 (bottom-right dashed box): These represent cases where the largest brain similarity is neither between the early-stage and later-stage MRI scans from the same fetuses nor between the later-stage FEM model and MRI scans from the same fetuses. Case 2, along with its extracted sulcal pits from the FEM scans and MRI scans, is included as an example of this category.

To further evaluate the M models’ performance, we selected the cases that demonstrated higher sulcal pits similarity between early-stage and later-stage MRI scans of the same fetuses, despite confounding effects of different longitudinal scan times across subjects. For these cases, we generated an SDSP heatmap for both early-stage and later-stage FEM models, as shown in Fig. 8. In Fig. 8, all diagonal values are higher than off-diagonal values within the same row, demonstrating that the sulcal pits from the later-stage FEM models (t_7_) align with their corresponding early-stage FEM models and the two FEM models from the same fetuses can be matched with 100% accuracy. This result suggests that the FEM models reliably preserve sulcal pit distribution during brain development. Additionally, the findings suggest that when accurate ground-truth data at the early stage is available, FEM models can effectively predict the spatiotemporal evolution of sulcal pits. The SDSP heatmap for all FEM models, comparing their early and later stages, is shown in Fig. S1. In total, five cases (cases 2, 4, 6, 12, and 13) in Fig. S1 were matched to later-stage FEM models of different cases (cases 11, 24, 24, 24, 24), failing to exhibit a higher SDSP between their associated early-stage and later-stage FEM models. A detailed inspection revealed that the initial scan times for three failed cases (cases 2, 4, and 13) had relatively smaller GWs, making the fetal brains appear less similar to their own FEM models (Fig. 5). Another observation is that the matched cases (cases 11 and 24) have smaller GWs at the later-stage scans. These are consistent with the explanation discussed in Fig. 5. Therefore, for more accurate prediction of the brain’s folded morphology based on its initial state, it is essential that the initial morphology reflects a time when primary folds have already formed on the brain’s surface. This also suggests that human brains at early stages of development share greater similarities and, with gradual growth, develop distinct folding patterns and sulcal pits.

**Fig. 8.**
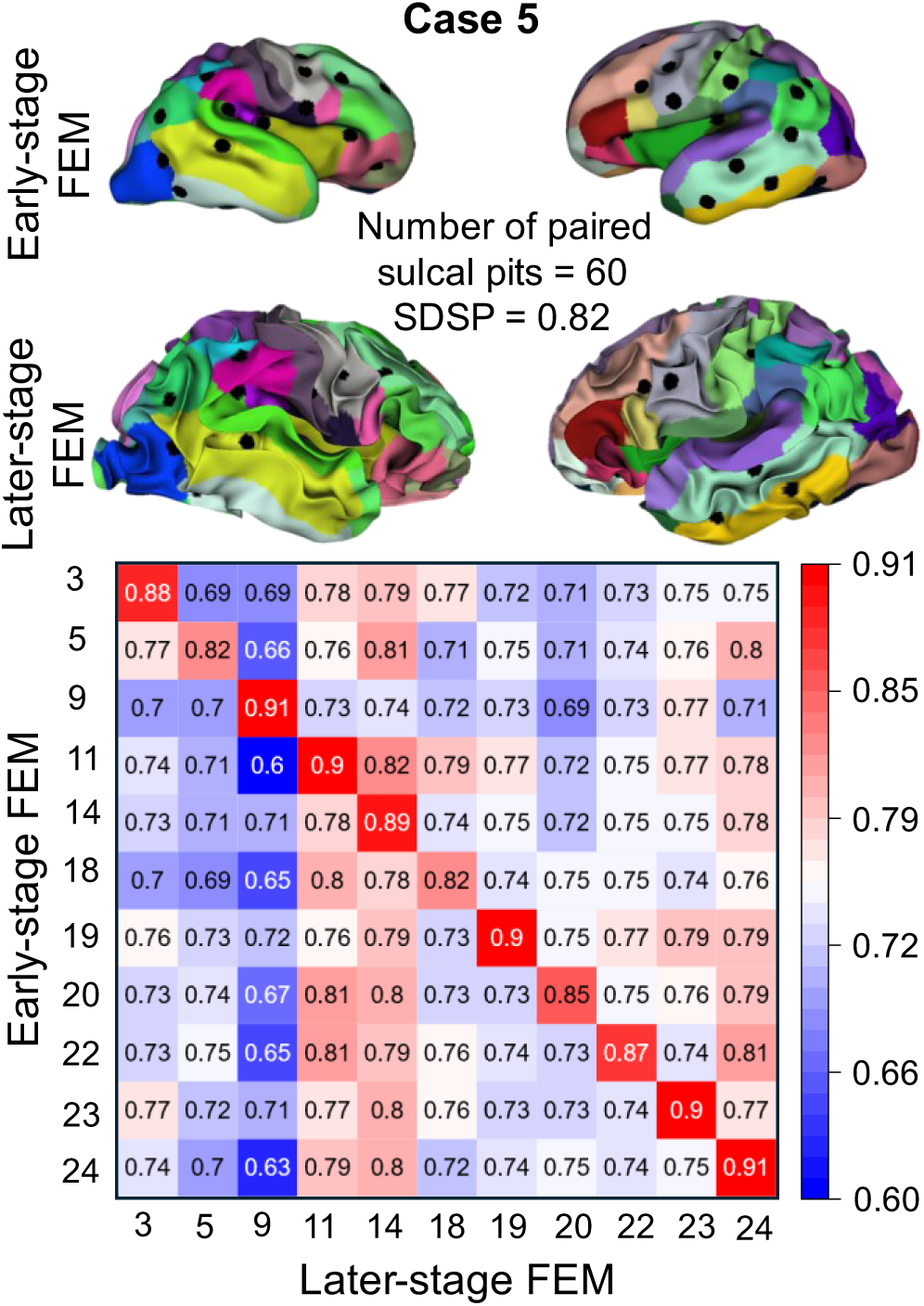
Sulcal pits-based similarity (SDSP) between the early-stage and later-stage FEM models for subjects with higher diagonal values in both ground truth and FEM-MRI analysis. The diagonal values represent within-subject similarity, while the off-diagonal values indicate between-subject similarity. Case 5, along with its extracted sulcal pits from the early- and later-stage FEM models, is presented as an example.

### Evolution and establishment of sulcal pits

A major challenge in understanding the spatiotemporal evolution of sulcal pits in individual brains is the limited availability of longitudinal data for each case. To examine how FEM models can address this limitation, we extracted all sulcal pits at each stage of the FEM models (from t_0_ to t_7_), matched them with the corresponding sulcal pits in early- and later-stage MRIs, and aligned to their respective gestational weeks. Figure 9a illustrates the relationship between gestational weeks and the evolution of the number of sulcal pits. As gestational weeks progress, the number of sulcal pits increases, reflecting the brain’s growing complexity in folding and fissure development. This figure demonstrates that FEM models serve as a valuable tool for bridging the gap between two longitudinal datasets. Notably, the figure reveals two distinct stages in the evolution of sulcal pits. During the initial stage, the number of sulcal pits in the FEM models increases in a relatively linear manner. In the second stage, however, the number of sulcal pits stabilizes, with minimal change observed. This transition highlights a key point in cortical development when folding becomes more established, aligning with observed patterns in neuroimaging studies^13^. This two-stage development of sulcal pits is evident in the MRI data shown in Fig. 9a. In the early gestational weeks, the number of sulcal pits increases rapidly, whereas in the later stages, their numbers stabilize with minimal change. This clear alignment between the FEM models and MRI data underscores the effectiveness of the FEM models in predicting brain folding and the spatiotemporal evolution of sulcal pits, providing a robust tool for understanding cortical development. Table S1 presents the number of sulcal pits for all 24 cases at each growth stage (t_1_ to t_7_), as well as the number of sulcal pits in the corresponding MRI scans at the early and later stages. Figure S2 represents the distribution of sulcal pits across the early and later stages of eight FEM models.

**Fig. 9.**
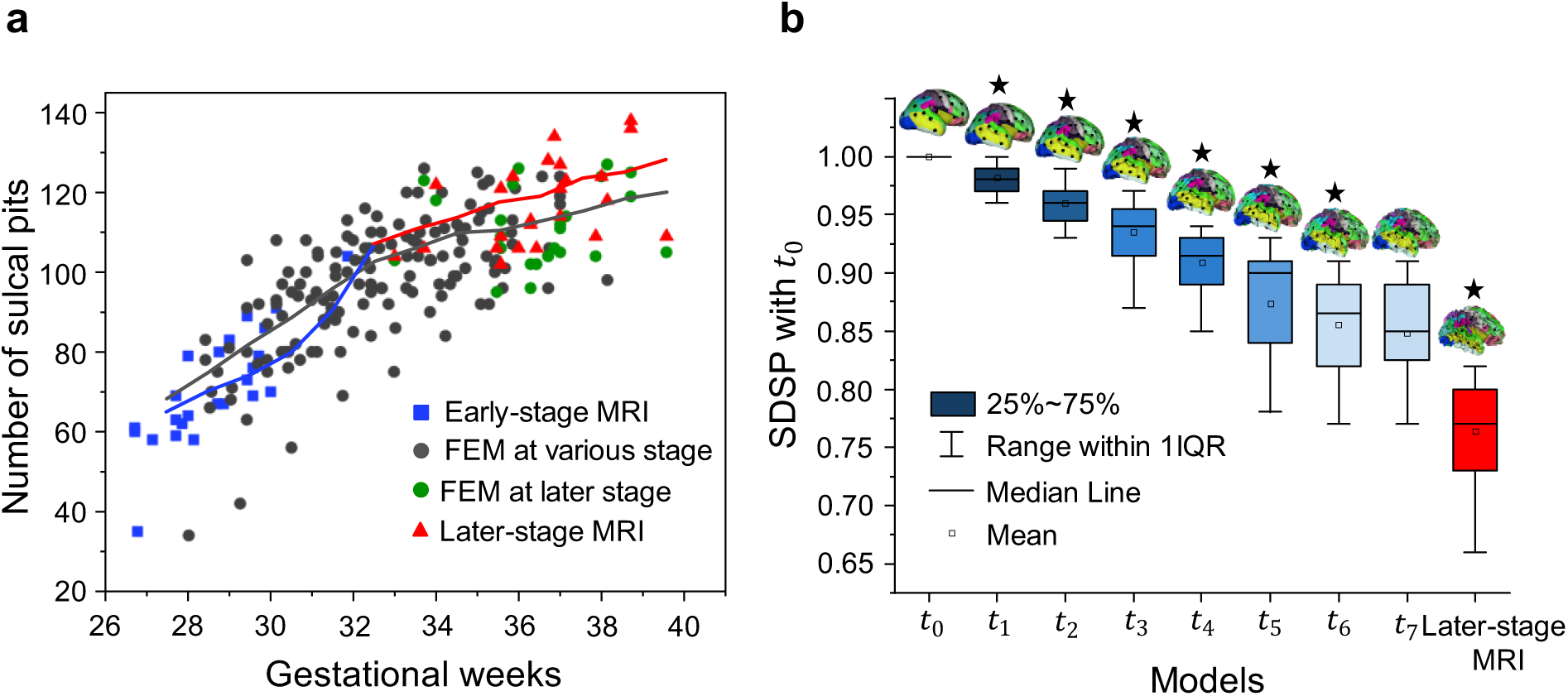
a. Relationship between gestational weeks and the number of sulcal pits. Squares represent early-stage MRI, and triangles denote later-stage MRI, with models from the same subject marked in identical colors. Black dots indicate FEM models sulcal pits across various developmental stages. The black line represents the moving average of FEM models, calculated using a window size of 3 and step size of 1; the blue and red lines show the moving averages of early and later-stage MRI scans, respectively. **b** Similarity degree of sulcal pits (SDSP) between early-stage MRI and various M models’ stages/later-stage MRI. The leftmost bar represents the SDSP of early-stage MRI with itself (value of 1), blue bars show SDSP between early-stage MRI and M models’ stages, and the red bar (rightmost) indicates SDSP between early and later-stage MRI. Data includes all 24 cases, with error bars indicating standard deviation. Each bar is topped with an example brain model showing sulcal pits (black dots). An asterisk above an example brain model denotes statistically significant difference between two bars at t_i v_s. t_i-1 (_i = 1, …, 7). The bar at the later-stage MRI was compared with the bar at t_7._

To investigate how sulcal pits establish in the brain model, we extracted the SDSP between all the simulated developmental stages (t_1_, t_2_, …, t_7_) and the early-stage MRI brain (t_0_) of each fetal brain. As the fetal brain develops from t_1_ to t_7_, we observed a decrease in brain similarity of sulcal pits with t_0_ (Fig. 9b). The SDSP of two brain models between each developmental stage (t_1_, t_2_, …, t_7_) and t_0_ is significantly lower (p < 0.001) than SDSP of two brain models between its previous stage (t_0_, t_1_, …, t_6_) and t_0_. However, there is no significant difference between SDSP of t_7_ and t_0_ and SDSP of t_6_ and t_0_ (p = 0.21). We also observed that the SDSP of the brain models between t_7_ and t_0_ was significantly higher than that between the later-stage MRI brain model and early-stage MRI scan (at t_0_). This discrepancy may be attributed to segmentation issues arising from the compromised quality of the fetal brain MRI scans at the later stages, possibly due to the compromised quality of the fetal brain MRI scans at the later stages, possibly due to fetal movement. On the other hand, this highlights the advantage of FEM models in reliably predicting the evolution of sulcal pits in the brain. An interesting observation in Fig. 9b is that once the folds and fissures are established in the model (between t_6_ and t_7_), there is minimal change in both the number and locations of the sulcal pits. This indicates their stability during brain growth and folding, a pattern consistent with imaging observations.

## Discussion

In this study, we developed true-scale, image-based mechanical models to elucidate how brain folds form during development and how sulcal pits evolve and establish within the folded structure. Our mechanical model demonstrates that the faster-growing cortical layer overlying the more slowly growing white matter core induces compressive stresses in the cortex, leading to invagination of the cortex into the white matter and the formation of folds^59^. As growth progresses, these folds become increasingly complex, giving rise to secondary and tertiary folds that resemble those observed in the mature brain. The simulations reveal that initial surface undulations (primary folds) during early developmental stages serve as critical locations guiding the formation of later-developing gyri and sulci. This finding aligns with other modeling studies that, even with simplified geometries, suggest fold positioning is strongly influenced by initial surface undulations^56^. This effect is especially pronounced in the formation and establishment of sulcal pits, as these are the deepest sulcal roots that remain preserved in position despite substantial cortical growth and brain size increase^17^.

The results of this study indicate that our models are powerful tools for dynamically capturing brain growth and folding processes, as well as for explaining the evolution and conservation of sulcal pits. A primary advantage of these mechanical models lies in their ability to trace the progression of surface morphology and sulcal pits across an arbitrary number of time points, a capacity limited in imaging studies due to the scarcity of longitudinal scans from individual brains^13^. Thus, these simulations serve as a valuable complement to imaging data, enhancing our understanding of the brain’s dynamic developmental trajectory. The results in Fig. 6 show that models derived from high-quality imaging data can accurately capture brain growth, folding, and the evolving patterning of sulcal pits. With reliable early-stage fetal brain inputs, the model can predict, with high accuracy, subsequent surface morphology and sulcal pit patterns post-growth and folding. This predictive capability represents a significant advancement in forecasting patient-specific sulcal pit development, especially since sulcal pit patterns are highly sensitive markers for detecting subtle abnormalities that traditional cortical measures, such as gyrification index and curvature, may overlook^13^.

The simulations indicate that sulcal pits act as anchor points in the folding morphology of the brain. During the brain’s expansion and folding, the number of sulcal pits increases, while previously formed pits remain stable, serving as conserved landmarks. Notably, as shown in Fig. 9, when the brain is highly folded and coincides with tertiary folding, the number of sulcal pits exhibits only a slight increase, reflecting the establishment of these features. Importantly, these pits are conserved even after birth and throughout infancy^17^. Consequently, sulcal pits can serve as a valuable proxy for linking prenatal and postnatal periods, while simultaneously addressing the heterogeneity of folding morphologies across individual brains. The models presented in this study can be further extended to replicate postnatal periods, supporting imaging studies that have demonstrated the position and spatial variance of sulcal pits in the fetal brain are similar to those in the adult brain^19^. In line with these observations, our mechanical models corroborate imaging findings regarding the establishment of sulcal pits in the fetal brain, particularly during the rapid growth of the third trimester, which underscores the stable locations of the earliest cortical sulci. urthermore, the model’s predictions regarding the increasing number of sulcal pits align with in vivo fetal MRI observations, indicating that sulcal emergence occurs during specific time frames in fetal brain development. Together, these findings enhance our understanding of how sulcal pits formation influences brain morphology across developmental stages^91–94^.

One interesting observation from Fig. 7 is that, in certain instances where the early-stage and later-stage MRI scans cannot be matched to their own cases based on SDSP, while the FEM model demonstrates its capability in simulating the folding process and preserving sulcal pits. This is evidenced by the higher similarity between the later-stage FEM model and the corresponding MRI scan from the same fetuses. Such findings suggest that the later-stage brain model generated using FEM exhibits a more comparable distribution of sulcal pits with the later-stage MRI scan from the same fetus. The later-stage FEM model can mimic the progression of sulcal pits throughout growth and folding although these results are still confounded by geometric imperfections present in fetal MRI scans. This highlights the robustness of computational modeling in accurately capturing sulcal pits distributions, even in cases where imaging data face quality challenges.

Recently, the accurate timing of sulcal emergence has been utilized as a reliable index for estimating gestational age and detecting developmental disorders^95^. Abnormal patterns of sulcal pits have been employed to study several brain disorders, including Down Syndrome^96^, polymicrogyria^97^, “isolated” agenesis of corpus callosum^98^, congenital heart disease^99^, and ASD^100^. Consequently, predictive models show great promise in simulating the development of the brain in relation to these specific disorders. This represents a significant step forward in understanding the mechanics of brain disorders that have often been overlooked by the brain mechanics community. In fact, these models offer the opportunity to manipulate various parameters, thereby elucidating their effects on both normal and disordered brain development. Such possibilities are not feasible with imaging data, which do not allow for the manipulation of growth and geometric parameters to study their contributions to brain development. Therefore, computational models serve as powerful tools for understanding the cause-and-effect relationships underlying brain development and disorders, ultimately enhancing our ability to identify and address neurodevelopmental challenges.

In our future studies, we aim to gain a deeper understanding of the mechanisms underlying brain disorders characterized by atypical development of the spatiotemporal evolution of sulcal pits. Our long-term hypothesis posits that abnormal patterns of sulcal pits observed after birth originate from atypical development during the fetal stage. One potential candidate for this investigation is ASD, which manifests atypical sulcal pit patterning during infancy and later stages^100,101^. Given that ASD symptoms typically emerge after birth, primarily during infancy and childhood, there is limited knowledge regarding its development during the fetal stage. Nevertheless, based on the atypical establishment of sulcal pits observed postnatally in ASD, we speculate that the disorder may also develop abnormally during the fetal period. Consequently, modeling sulcal pits could serve as a proxy to link prenatal and postnatal stages of development. Investigating the development of sulcal pits in ASD, a heterogeneous disorder primarily attributed to genetic factors, may help clarify the numerous contradictory findings related to alterations in cortical layer thickness, surface area, volume, and gyrification index associated with ASD^102–106^. Therefore, we propose that examining the mechanics of typical sulcal pits development provides a valuable foundation for further studies on the development of ASD.

Finally, akin to other modeling studies, our research acknowledges certain simplifications and limitations inherent in computational models. In this study, we focused exclusively on differential tangential growth, which is a complex phenomenon that may arise from various underlying processes. However, other significant contributing factors, such as the impact of neural connectivity and its role in cortical development, were not incorporated into our model^52,55,57,107,108^. The material properties utilized in our simulations were assumed to be constant. Yet, existing literature indicates that the mechanical properties of both the cortical plate and white matter can evolve throughout development, affecting the overall mechanical behavior of the brain^109^. This variability underscores the importance of integrating time-dependent material properties in future modeling efforts to enhance accuracy. Furthermore, our current models did not account for hemispheric asymmetries or sex differences, both of which have been shown to influence the morphology of sulcal pits in fetal brains^95^. These factors could contribute to variability in the development of brain structures, indicating that future studies should explicitly incorporate these parameters to facilitate a more nuanced discussion of their effects. In this study, sulcal pits were extracted using the concept of sulcal depth; however, previous research has also employed mean curvature as an extraction method. While both approaches are applicable to fetal brain analysis and yield comparable results, discrepancies persist between the two methods^19^. Establishing a reliable geometric feature for sulcal pits extraction could provide a clearer understanding of these differences and improve the robustness of our findings. Ultimately, addressing these limitations will strengthen future modeling studies and enhance our ability to explore the complexities of brain development.

## Conclusion

In this study, we developed and evaluated the first true-scale, image-based mechanical model to explore the spatiotemporal evolution of brain sulcal pits in individual fetal brains. Constructed using scans from the initial time point of longitudinal data, our model predicts the brain’s surface morphology by comparing sulcal pits between model predictions and later time point scans. This dynamic approach elucidates the transformation of a smooth fetal brain with primary folds into one characterized by secondary and tertiary folds. Our findings align with imaging data, demonstrating that sulcal pits remain stable during brain development and can serve as crucial markers linking prenatal and postnatal brain characteristics. The main advantage of the mechanical models lies in their ability to trace the progression of surface morphology and sulcal pits across an arbitrary number of time points, a capacity limited in imaging studies due to the scarcity of longitudinal scans from individual brains. The developed model provides a robust platform for studying the evolution of sulcal pits in both healthy and disordered brains, which is particularly significant given that altered patterns of sulcal pits are observed in disorders. Ultimately, our study lays a solid foundation for future investigations into the complex mechanisms driving brain morphology and their implications for understanding neurodevelopmental challenges.

## Acknowledgment

This work was supported by the National Science Foundation under grant number CMMI: 2123061, awarded to M. J. Razavi, W. Dai, and A. Gholipour, and by the National Institutes of Health under award number R01EB031849, awarded to A. Gholipour.

## Author contributions

A. S.: Methodology, Formal analysis, Software, Writing – original draft.

1. Y. G.: Methodology, Formal analysis, Software, Writing – original draft.

A. G.: Formal analysis, Writing – review & editing.

W.D.: Investigation, Supervision, Writing – review & editing, Funding acquisition.

M. J. R.: Conceptualization, Investigation, Supervision, Writing – review & editing, Funding acquisition. All authors reviewed the manuscript.

## Data and code availability

The data and code underlying the findings of this study will be made available through the Zenodo repository upon publication

## Supplementary Information

**Fig. S1.**
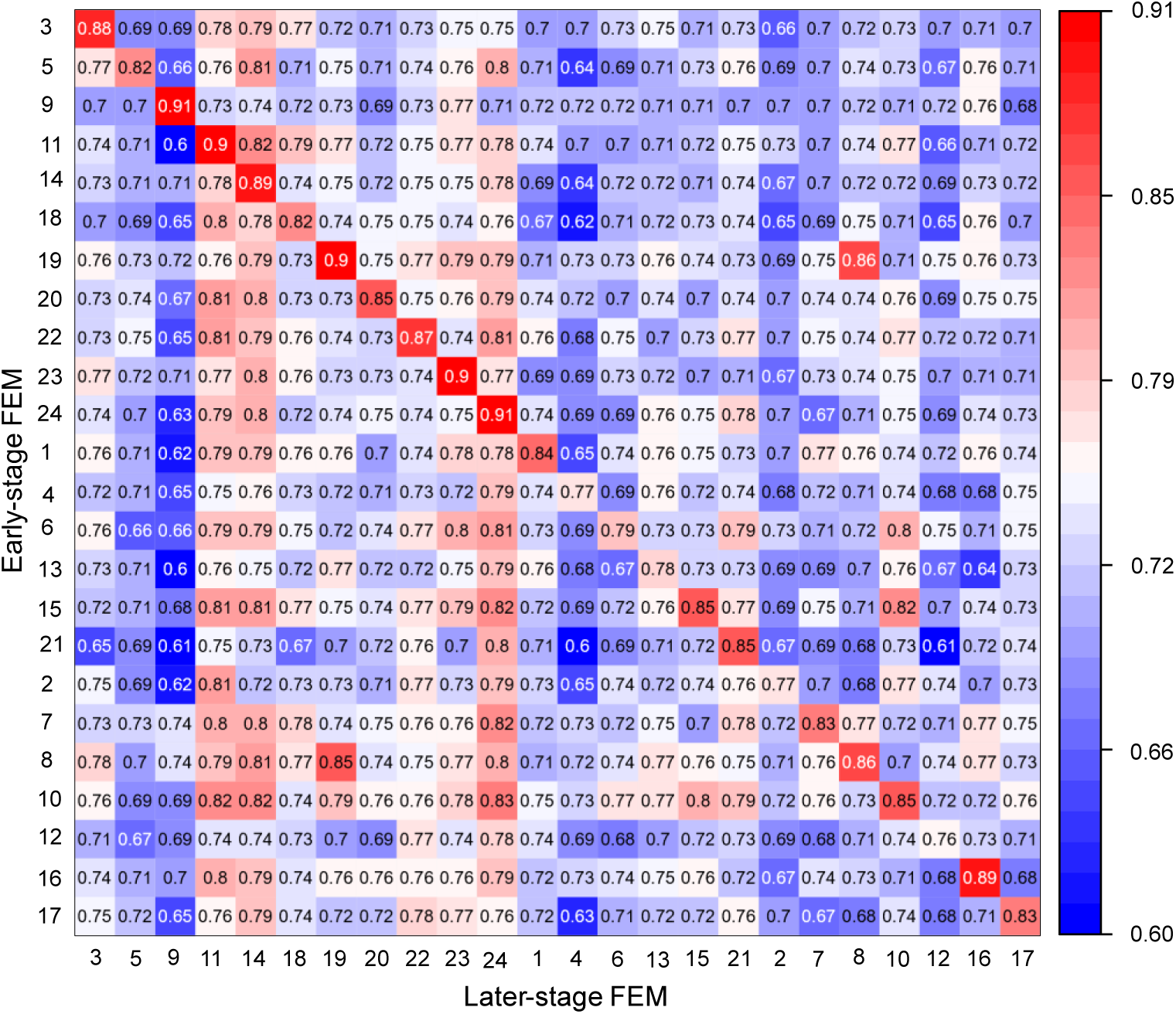
Sulcal pits-based similarity (SDSP) between early-stage FEM and later-stage FEM for all subject pairs. The diagonal indicates similarity within the same subject, while the off diagonal reflects similarity

**Table S1.** Number of sulcal pits for all 24 cases at each growth stage (t_1 t_o t_7)_, as well as the number of sulcal pits in the corresponding MRI scans at the early and later stages.

| Case number | $t_0$<br>MRI early-stage<br>(same for FEM) | $t_1$ | $t_2$ | $t_3$ | $t_4$ | $t_5$ | $t_6$ | $t_7$ | MRI Later-stage |
| --- | --- | --- | --- | --- | --- | --- | --- | --- | --- |
| 1 | 91 | 93 | 97 | 104 | 111 | 113 | 117 | 127 | 118 |
| 2 | 58 | 66 | 78 | 87 | 96 | 97 | 103 | 106 | 134 |
| 3 | 89 | 95 | 105 | 110 | 113 | 117 | 121 | 111 | 127 |
| 4 | 61 | 83 | 108 | 113 | 120 | 123 | 124 | 125 | 136 |
| 5 | 70 | 80 | 92 | 99 | 99 | 96 | 106 | 104 | 109 |
| 6 | 58 | 63 | 78 | 85 | 96 | 105 | 107 | 114 | 123 |
| 7 | 67 | 75 | 93 | 103 | 112 | 104 | 104 | 102 | 113 |
| 8 | 69 | 76 | 88 | 97 | 103 | 103 | 101 | 106 | 109 |
| 9 | 104 | 106 | 107 | 108 | 110 | 112 | 113 | 113 | 121 |
| 10 | 35 | 34 | 42 | 56 | 69 | 75 | 84 | 95 | 106 |
| 11 | 64 | 75 | 103 | 92 | 100 | 101 | 103 | 103 | 104 |
| 12 | 79 | 91 | 108 | 116 | 126 | 122 | 124 | 124 | 124 |
| 13 | 60 | 78 | 93 | 100 | 107 | 116 | 119 | 119 | 138 |
| 14 | 86 | 97 | 99 | 110 | 120 | 120 | 125 | 122 | 124 |
| 15 | 62 | 71 | 80 | 91 | 96 | 103 | 92 | 96 | 112 |
| 16 | 79 | 82 | 94 | 101 | 104 | 111 | 115 | 126 | 106 |
| 17 | 63 | 68 | 80 | 80 | 86 | 97 | 101 | 105 | 114 |
| 18 | 80 | 87 | 95 | 106 | 108 | 108 | 110 | 112 | 121 |
| 19 | 76 | 80 | 84 | 90 | 92 | 96 | 98 | 105 | 109 |
| 20 | 73 | 80 | 89 | 95 | 95 | 99 | 102 | 102 | 106 |
| 21 | 69 | 81 | 89 | 88 | 92 | 94 | 100 | 104 | 128 |
| 22 | 67 | 77 | 94 | 90 | 106 | 110 | 110 | 109 | 102 |
| 23 | 83 | 92 | 100 | 104 | 109 | 114 | 109 | 118 | 122 |
| 24 | 59 | 70 | 80 | 97 | 105 | 108 | 193 | 123 | 106 |

**Fig. S2.**
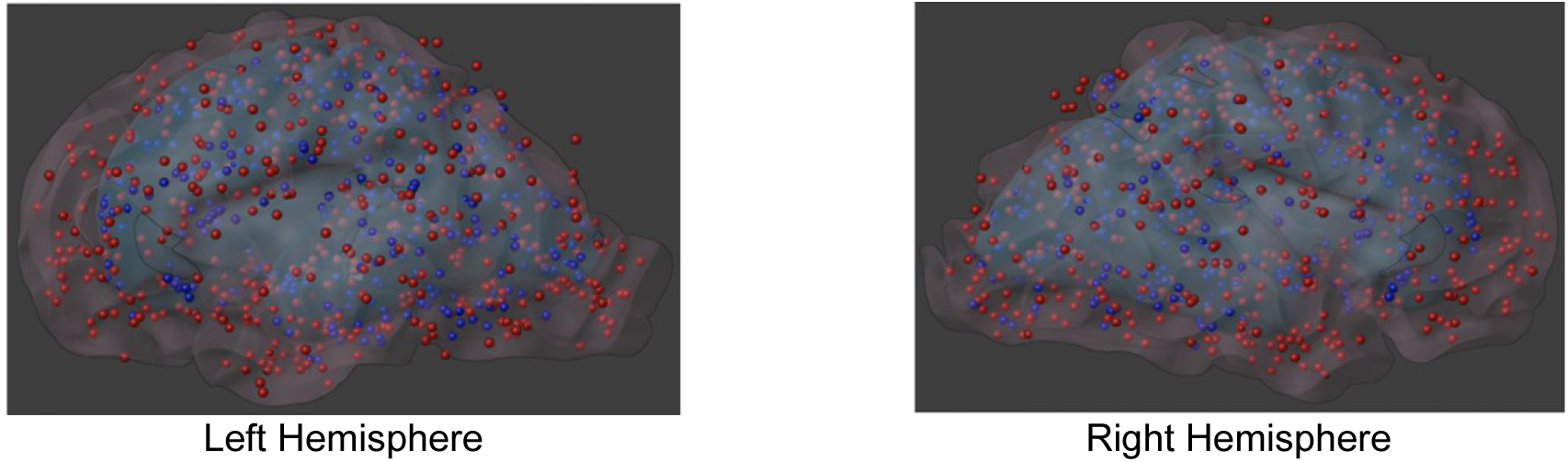
Distribution of sulcal pits across the early and later stages of eight brain growth models. The red surface represents the later-stage FEM, while the blue surface corresponds to the early-stage FEM. Red dots denote the sulcal pits at the later stage, while blue dots indicate the corresponding sulcal pits at the early stage.

## References

1. Zilles, K., Palomero-Gallagher, N. & Amunts, K. Development of cortical folding during evolution and ontogeny. Trends Neurosci. 36, 275–284 (2013).

2. Striedter, G. F., Srinivasan, S. & Monuki, E. S. Cortical Folding: When, Where, How, and Why? Annu. Rev. Neurosci. 38, 291–307 (2015).

3. Fernández, V., Llinares-Benadero, C. & Borrell, V. Cerebral cortex expansion and folding: what have we learned? EMBO J. 35, 1021–1044 (2016).

4. Welker, W. Why Does Cerebral Cortex Fissure and Fold? in Cerebral Cortex (eds. Jones, E. G. & Peters, A.) vol. 8B 3–136 (Springer US, Boston, MA, 1990).

5. Ten Donkelaar, H. J. et al. Overview of the Development of the Human Brain and Spinal Cord. in Clinical Neuroembryology 1–76 (Springer International Publishing, Cham, 2023). doi:10.1007/978-3-031-26098-8_1.

6. Del-Valle-Anton, L. & Borrell, V. Folding brains: from development to disease modeling. Physiol. Rev. 102, 511–550 (2022).

7. Sun, T. & Hevner, R. F. Growth and folding of the mammalian cerebral cortex: from molecules to malformations. Nat. Rev. Neurosci. 15, 217–232 (2014).

8. Hardan, A. Y., Jou, R. J., Keshavan, M. S., Varma, R. & Minshew, N. J. Increased frontal cortical folding in autism: a preliminary MRI study. Psychiatry Res. Neuroimaging 131, 263– 268 (2004).

9. Harris, J. M. et al. Abnormal cortical folding in high-risk individuals: a predictor of the development of schizophrenia? Biol. Psychiatry 56, 182–189 (2004).

10. Choi, K. W. et al. Decreased cortical gyrification in patients with bipolar disorder. Psychol. Med. 52, 2232–2244 (2022).

11. Fernández, V. & Borrell, V. Developmental mechanisms of gyrification. Curr. Opin. Neurobiol. 80, 102711 (2023).

12. Ronan, L. & Fletcher, P. C. From genes to folds: a review of cortical gyrification theory. Brain Struct. Funct. 220, 2475–2483 (2015).

13. Im, K. & Grant, P. E. Sulcal pits and patterns in developing human brains. NeuroImage 185, 881–890 (2019).

14. Régis, J. et al. “Sulcal Root” eneric Model: a Hypothesis to Overcome the Variability of the Human Cortex Folding Patterns. Neurol. Med. Chir. (Tokyo) 45, 1–17 (2005).

15. Lohmann, G., von Cramon, D. Y. & Colchester, A. C. F. Deep Sulcal Landmarks Provide an Organizing Framework for Human Cortical Folding. Cereb. Cortex 18, 1415–1420 (2008).

16. Im, K. et al. Spatial Distribution of Deep Sulcal Landmarks and Hemispherical Asymmetry on the Cortical Surface. Cereb. Cortex 20, 602–611 (2010).

17. Meng, Y., Li, G., Lin, W., Gilmore, J. H. & Shen, D. Spatial distribution and longitudinal development of deep cortical sulcal landmarks in infants. NeuroImage 100, 206–218 (2014).

18. Dubois, J. et al. Mapping the Early Cortical Folding Process in the Preterm Newborn Brain. Cereb. Cortex 18, 1444–1454 (2008).

19. Yun, H. J. et al. Temporal Patterns of Emergence and Spatial Distribution of Sulcal Pits During Fetal Life. Cereb. Cortex 30, 4257–4268 (2020).

20. Zilles, K. et al. Quantitative analysis of sulci in the human cerebral cortex: development, regional heterogeneity, gender difference, asymmetry, intersubject variability and cortical architecture. Hum. Brain Mapp. 5, 218–221 (1997).

21. Hasnain, M. K. Structure-Function Spatial Covariance in the Human Visual Cortex. Cereb. Cortex 11, 702–716 (2001).

22. Eickhoff, S. B., Heim, S., Zilles, K. & Amunts, K. Testing anatomically specified hypotheses in functional imaging using cytoarchitectonic maps. NeuroImage 32, 570–582 (2006).

23. Fischl, B. et al. Cortical Folding Patterns and Predicting Cytoarchitecture. Cereb. Cortex 18, 1973–1980 (2008).

24. Auzias, G., Brun, L., Deruelle, C. & Coulon, O. Deep sulcal landmarks: Algorithmic and conceptual improvements in the definition and extraction of sulcal pits. NeuroImage 111, 12–25 (2015).

25. Fukuchi-Shimogori, T. & Grove, E. A. Neocortex Patterning by the Secreted Signaling Molecule FGF8. Science 294, 1071–1074 (2001).

26. Miyashita-Lin, E. M., Hevner, R., Wassarman, K. M., Martinez, S. & Rubenstein, J. L. R. Early Neocortical Regionalization in the Absence of Thalamic Innervation. Science 285, 906–909 (1999).

27. Rakic, P. Specification of cerebral cortical areas. Science 241, 170–176 (1988).

28. Rakic, P. Neurocreationism--Making New Cortical Maps. Science 294, 1011–1012 (2001).

29. Le Guen, Y. et al. Genetic Influence on the Sulcal Pits: On the Origin of the First Cortical Folds. Cereb. Cortex 28, 1922–1933 (2018).

30. Im, K. et al. Quantitative comparison and analysis of sulcal patterns using sulcal graph matching: A twin study. NeuroImage 57, 1077–1086 (2011).

31. Im, K. et al. Quantitative Folding Pattern Analysis of Early Primary Sulci in Human Fetuses with Brain Abnormalities. *Am*. J. Neuroradiol. 38, 1449–1455 (2017).

32. Huang, Y. et al. Genetic Influence on Gyral Peaks. NeuroImage 280, 120344 (2023).

33. Bayly, P. V., Taber, L. A. & Kroenke, C. D. Mechanical forces in cerebral cortical folding: A review of measurements and models. J. Mech. Behav. Biomed. Mater. 29, 568–581 (2014).

34. Budday, S., Steinmann, P. & Kuhl, E. The role of mechanics during brain development. J. Mech. Phys. Solids 72, 75–92 (2014).

35. Razavi, M. J., Zhang, T., Li, X., Liu, T. & Wang, X. Role of mechanical factors in cortical folding development. *Phys*. Rev. E 92, 032701 (2015).

36. Hilgetag, C. C. & Barbas, H. Role of Mechanical Factors in the Morphology of the Primate Cerebral Cortex. PLoS Comput. Biol. 2, e22 (2006).

37. de Vareilles, H., Rivière, D., Mangin, J. F. & Dubois, J. Development of cortical folds in the human brain: An attempt to review biological hypotheses, early neuroimaging investigations and functional correlates. Dev. Cogn. Neurosci. 61, 101249 (2023).

38. Garcia, K. E., Kroenke, C. D. & Bayly, P. V. Mechanics of cortical folding: stress, growth and stability. Philos. Trans. R. Soc. B Biol. Sci. 373, 20170321 (2018).

39. Caviness, V. Mechanical model of brain convolutional development. Science 189, 18–21 (1975).

40. Tallinen, T., Chung, J. Y., Biggins, J. S. & Mahadevan, L. Gyrification from constrained cortical expansion. Proc. Natl. Acad. Sci. 111, 12667–12672 (2014).

41. Tallinen, T. et al. On the growth and form of cortical convolutions. Nat. Phys. 12, 588–593 (2016).

42. Razavi, M. J., Zhang, T., Liu, T. & Wang, X. Cortical Folding Pattern and its Consistency Induced by Biological Growth. Sci. Rep. 5, (2015).

43. Bayly, P. V., Okamoto, R. J., Xu, G., Shi, Y. & Taber, L. A. A cortical folding model incorporating stress-dependent growth explains gyral wavelengths and stress patterns in the developing brain. Phys. Biol. 10, 016005 (2013).

44. Balouchzadeh, R., Bayly, P. V. & Garcia, K. E. Effects of stress-dependent growth on evolution of sulcal direction and curvature in models of cortical folding. Brain Multiphysics 4, 100065 (2023).

45. Budday, S., Raybaud, C. & Kuhl, E. A mechanical model predicts morphological abnormalities in the developing human brain. Sci. Rep. 4, 5644 (2015).

46. Zhang, T. et al. Mechanisms of circumferential gyral convolution in primate brains. J. Comput. Neurosci. 42, 217–229 (2017).

47. Wang, S., Saito, K., Kawasaki, H. & Holland, M. A. Orchestrated neuronal migration and cortical folding: A computational and experimental study. PLOS Comput. Biol. 18, e1010190 (2022).

48. Wang, S., Demirci, N. & Holland, M. A. Numerical investigation of biomechanically coupled growth in cortical folding. Biomech. Model. Mechanobiol. 20, 555–567 (2021).

49. Darayi, M. et al. Computational models of cortical folding: A review of common approaches. J. Biomech. 139, 110851 (2022).

50. Rakic, P. A small step for the cell, a giant leap for mankind: a hypothesis of neocortical expansion during evolution. Trends Neurosci. 18, 383–388 (1995).

51. Chen, H. et al. A Dynamic Skull Model for Simulation of Cerebral Cortex Folding. in Medical Image Computing and Computer-Assisted Intervention – MICCAI 2010 (eds. Jiang, T., Navab, N., Pluim, J. P. W. & Viergever, M. A.) vol. 6362 412–419 (Springer Berlin Heidelberg, Berlin, Heidelberg, 2010).

52. Essen, D. C. V. A tension-based theory of morphogenesis and compact wiring in the central nervous system. Nature 385, 313–318 (1997).

53. Van Essen, D. C. A 2020 view of tension-based cortical morphogenesis. Proc. Natl. Acad. Sci. 117, 32868–32879 (2020).

54. Garcia, K. E., Wang, X. & Kroenke, C. D. A model of tension-induced fiber growth predicts white matter organization during brain folding. Nat. Commun. 12, 6681 (2021).

55. Chavoshnejad, P. et al. Role of axonal fibers in the cortical folding patterns: A tale of variability and regularity. Brain Multiphysics 2, 100029 (2021).

56. Chavoshnejad, P. et al. Mechanical hierarchy in the formation and modulation of cortical folding patterns. Sci. Rep. 13, 13177 (2023).

57. Holland, M. A., Miller, K. E. & Kuhl, E. Emerging Brain Morphologies from Axonal Elongation. Ann. Biomed. Eng. 43, 1640–1653 (2015).

58. Razavi, M. J., Zhang, T., Li, X., Liu, T. & Wang, X. Role of mechanical factors in cortical folding development. *Phys*. Rev. E 92, (2015).

59. Tallinen, T. et al. On the growth and form of cortical convolutions. Nat. Phys. 12, 588–593 (2016).

60. Bayly, P. V., Okamoto, R. J., Xu, G., Shi, Y. & Taber, L. A. A cortical folding model incorporating stress-dependent growth explains gyral wavelengths and stress patterns in the developing brain. Phys. Biol. 10, 016005 (2013).

61. Toro, R. & Burnod, Y. A Morphogenetic Model for the Development of Cortical Convolutions. Cereb. Cortex 15, 1900–1913 (2005).

62. Budday, S., Steinmann, P., Goriely, A. & Kuhl, E. Size and curvature regulate pattern selection in the mammalian brain. Extreme Mech. Lett. 4, 193–198 (2015).

63. Campos, L. da C., Hornung, R., Gompper, G., Elgeti, J. & Caspers, S. The role of thickness inhomogeneities in hierarchical cortical folding. ArXiv200401020 Cond-Mat Physicsphysics Q-Bio (2020).

64. Leyva-Mendivil, M. F., Page, A., Bressloff, N. W. & Limbert, G. A mechanistic insight into the mechanical role of the stratum corneum during stretching and compression of the skin. J. Mech. Behav. Biomed. Mater. 49, 197–219 (2015).

65. Wang, L. et al. A three-layer mechanical model for the analysis of effects of pia matter on cortical folding. Eng. Comput. 36, 2634–2650 (2019).

66. Wang, S., Demirci, N. & Holland, M. A. Numerical investigation of biomechanically coupled growth in cortical folding. Biomech. Model. Mechanobiol. (2020) doi:10.1007/s10237-020-01400-w.

67. Jafarabadi, F., Wang, S. & Holland, M. A. A Numerical Study on the Influence of Cerebrospinal Fluid Pressure on Brain Folding. J. Appl. Mech. 90, 071006 (2023).

68. Gholipour, A., Estroff, J. A. & Warfield, S. K. Robust Super-Resolution Volume Reconstruction From Slice Acquisitions: Application to Fetal Brain MRI. IEEE Trans. Med. Imaging 29, 1739–1758 (2010).

69. Kuklisova-Murgasova, M., Quaghebeur, G., Rutherford, M. A., Hajnal, J. V. & Schnabel, J. A. Reconstruction of fetal brain MRI with intensity matching and complete outlier removal. Med. Image Anal. 16, 1550–1564 (2012).

70. Kainz, B. et al. Fast Volume Reconstruction From Motion Corrupted Stacks of 2D Slices. IEEE Trans. Med. Imaging 34, 1901–1913 (2015).

71. Ebner, M. et al. An automated framework for localization, segmentation and super-resolution reconstruction of fetal brain MRI. NeuroImage 206, 116324 (2020).

72. Uus, A. et al. Deformable Slice-to-Volume Registration for Motion Correction of Fetal Body and Placenta MRI. IEEE Trans. Med. Imaging 39, 2750–2759 (2020).

73. Gholipour, A. et al. A normative spatiotemporal MRI atlas of the fetal brain for automatic segmentation and analysis of early brain growth. Sci. Rep. 7, 476 (2017).

74. Khan, S. et al. Fetal brain growth portrayed by a spatiotemporal diffusion tensor MRI atlas computed from in utero images. NeuroImage 185, 593–608 (2019).

75. Dou, H. et al. A Deep Attentive Convolutional Neural Network for Automatic Cortical Plate Segmentation in Fetal MRI. IEEE Trans. Med. Imaging 40, 1123–1133 (2021).

76. Karimi, D., Rollins, C. K., Velasco-Annis, C., Ouaalam, A. & Gholipour, A. Learning to segment fetal brain tissue from noisy annotations. Med. Image Anal. 85, 102731 (2023).

77. Payette, K. et al. Fetal brain tissue annotation and segmentation challenge results. Med. Image Anal. 88, 102833 (2023).

78. Dahnke, R. & Gaser, C. Surface and Shape Analysis. in Brain Morphometry (eds. Spalletta, G., Piras, F. & Gili, T.) vol. 136 51–73 (Springer New York, New York, NY, 2018).

79. Demirci, N. & Holland, M. A. Cortical thickness systematically varies with curvature and depth in healthy human brains. Hum. Brain Mapp. 43, 2064–2084 (2022).

80. Xue, H. et al. Automatic segmentation and reconstruction of the cortex from neonatal MRI. NeuroImage 38, 461–477 (2007).

81. Rodriguez, E. K., Hoger, A. & McCulloch, A. D. Stress-dependent finite growth in soft elastic tissues. J. Biomech. 27, 455–467 (1994).

82. Liu, M. et al. Robust Cortical Thickness Morphometry of Neonatal Brain and Systematic Evaluation Using Multi-Site MRI Datasets. Front. Neurosci. 15, 650082 (2021).

83. Ronan, L. et al. Differential Tangential Expansion as a Mechanism for Cortical Gyrification. Cereb. Cortex 24, 2219–2228 (2014).

84. Budday, S., Ovaert, T. C., Holzapfel, G. A., Steinmann, P. & Kuhl, E. Fifty Shades of Brain: A Review on the Mechanical Testing and Modeling of Brain Tissue. Arch. Comput. Methods Eng. (2019) doi:10.1007/s11831-019-09352-w.

85. Budday, S. et al. Mechanical properties of gray and white matter brain tissue by indentation. J. Mech. Behav. Biomed. Mater. 46, 318–330 (2015).

86. Budday, S. et al. Mechanical characterization of human brain tissue. Acta Biomater. 48, 319–340 (2017).

87. Rettmann, M. E., Han, X., Xu, C. & Prince, J. L. Automated Sulcal Segmentation Using Watersheds on the Cortical Surface. NeuroImage 15, 329–344 (2002).

88. Boucher, M., Whitesides, S. & Evans, A. Depth potential function for folding pattern representation, registration and analysis. Med. Image Anal. 13, 203–214 (2009).

89. Hutchison, D. et al. Reweighted Random Walks for Graph Matching. in Computer Vision – ECCV 2010 (eds. Daniilidis, K., Maragos, P. & Paragios, N.) vol. 6315 492–505 (Springer Berlin Heidelberg, Berlin, Heidelberg, 2010).

90. Cashbaugh, J. & Kitts, C. Automatic Calculation of a Transformation Matrix Between Two Frames. IEEE Access 6, 9614–9622 (2018).

91. Chi, J. G., Dooling, E. C. & Gilles, F. H. Gyral development of the human brain. Ann. Neurol. 1, 86–93 (1977).

92. Garel, C. et al. Fetal cerebral cortex: normal gestational landmarks identified using prenatal MR imaging. AJNR Am. J. Neuroradiol. 22, 184–189 (2001).

93. Fogliarini, C. et al. Assessment of cortical maturation with prenatal MRI. Part I: normal cortical maturation. Eur. Radiol. 15, 1671–1685 (2005).

94. Habas, P. A. et al. Early Folding Patterns and Asymmetries of the Normal Human Brain Detected from in Utero MRI. Cereb. Cortex 22, 13–25 (2012).

95. Yun, H. J. et al. Quantification of sulcal emergence timing and its variability in early fetal life: Hemispheric asymmetry and sex difference. NeuroImage 263, 119629 (2022).

96. Yun, H. J. et al. Regional Alterations in Cortical Sulcal Depth in Living Fetuses with Down Syndrome. Cereb. Cortex 31, 757–767 (2021).

97. Im, K. et al. Quantification and Discrimination of Abnormal Sulcal Patterns in Polymicrogyria. Cereb. Cortex 23, 3007–3015 (2013).

98. Tarui, T. et al. Disorganized Patterns of Sulcal Position in Fetal Brains with Agenesis of Corpus Callosum. Cereb. Cortex 28, 3192–3203 (2018).

99. Ortinau, C. M. et al. Early-Emerging Sulcal Patterns Are Atypical in Fetuses with Congenital Heart Disease. Cereb. Cortex 29, 3605–3616 (2019).

100. Brun, L. et al. ocalized Misfolding ithin Broca’s Area as a Distinctive eature of Autistic Disorder. Biol. Psychiatry Cogn. Neurosci. Neuroimaging 1, 160–168 (2016).

101. Auzias, G. et al. Atypical sulcal anatomy in young children with autism spectrum disorder. NeuroImage Clin. 4, 593–603 (2014).

102. Mensen, V. T. et al. Development of cortical thickness and surface area in autism spectrum disorder. NeuroImage Clin. 13, 215–222 (2017).

103. Ohta, H. et al. Increased Surface Area, but not Cortical Thickness, in a Subset of Young Boys With Autism Spectrum Disorder: Cortical thickness in autism spectrum disorder. Autism Res. 9, 232–248 (2016).

104. Khundrakpam, B. S., Lewis, J. D., Kostopoulos, P., Carbonell, F. & Evans, A. C. Cortical Thickness Abnormalities in Autism Spectrum Disorders Through Late Childhood, Adolescence, and Adulthood: A Large-Scale MRI Study. Cereb. Cortex 27, 1721–1731 (2017).

105. Smith, E. et al. Cortical thickness change in autism during early childhood: CT in Early Childhood ASD. Hum. Brain Mapp. 37, 2616–2629 (2016).

106. Hardan, A. Y., Muddasani, S., Vemulapalli, M., Keshavan, M. S. & Minshew, N. J. An MRI Study of Increased Cortical Thickness in Autism. Am. J. Psychiatry 163, 1290–1292 (2006).

107. Wang, X., Wang, S. & Holland, M. A. Axonal tension contributes to consistent fold placement. Soft Matter 20, 3053–3065 (2024).

108. Solhtalab, A., Foroughi, A. H., Pierotich, L. & Razavi, M. J. Stress landscape of folding brain serves as a map for axonal pathfinding. Nat. Commun. 16, 1187 (2025).

109. Walter, C. et al. Multi-scale measurement of stiffness in the developing ferret brain. Sci. Rep. 13, 20583 (2023).

